# Single-cell spatially resolved transcriptomic characterization of the developing mouse cochlea

**DOI:** 10.64898/2026.01.21.700589

**Authors:** Philippe Jean, Sabrina Mechaussier, Amrit Singh-Estivalet, Céline Trébeau, Aurore Gaudin, Laura Barrio Cano, Andrea Lelli, Fabienne Wong Jun Tai, Sébastien Megharba, Sandrine Schmutz, Sarra Loulizi, Sophie Novault, David Hardy, Carolina Moraes-Cabe, Milena Hasan, Christine Petit, Raphael Etournay, Nicolas Michalski

## Abstract

The cochlea, the sensory organ of hearing, functions as a frequency analyzer, analogous to a musical instrument. During development, while the medio-lateral axis supports differentiation of sensory cells and their surrounding supporting cells, the longitudinal axis underlies frequency-dependent properties of the cochlea. The combination of these two gene expression gradients defines unique physiological attributes of each cell intimately linked to its position within the cochlea. To determine which cochlear cell-types have a transcriptomic signature sensitive to these two gradients and identify the underlying gene regulatory networks, we took advantage of the advent of spatial single cell transcriptomics methodologies. We therefore generated a spatial transcriptomic atlas reaching single cell resolution based on the Visium HD technique, a sequencing-based technology that employ arrays of spatially barcoded probes to capture RNA molecules unbiasedly from histological tissue sections. Spatial transcriptional changes during embryonic stages, E14 and E16, as well as during postnatal development, P1 and P8, were investigated. Based on this dataset, not only cell-type assignment in single cell RNA-seq experiments could be validated, but the classification for some cell-types could be refined. Gradients of gene expression along the medio-lateral and longitudinal axes in multiple cell-types together with their temporal dynamics across development were also uncovered. Altogether, this atlas paves the way for deciphering gene regulatory networks controlling gene expression as a function of position in the cochlear cell types, providing a valuable resource for the design of efficient, robust and safe gene therapy strategies.

## INTRODUCTION

The cochlea, the mammalian sensory organ for hearing, requires to encode sounds at high speed and extreme temporal precision to extract as much information as possible from sound stimuli (Palmer and Russell, 1986). To meet these functional demands, the auditory sensory organ has evolved into a mechanical system capable of decomposing sounds into distinct frequency components, thereby allowing parallel processing of the information conveyed by each frequency band (Müller et al., 2005). With its longitudinally varying mechanical attributes, the cochlea is spatially organized to detect the whole range of sound frequencies, a feature known as tonotopy (Békésy and Wever, 1960), specific of each species that adapted to the most relevant sounds in their environment. Therefore, the auditory organ is a highly complex and organized structure harboring a high diversity of cochlear cell-types to support the function of the auditory sensory hair cells (HCs) (Lim, 1986). Moreover, being physically constrained into a bony structure; the mouse cochlea only harbors around 3 000 HCs in the mouse compared to millions of photoreceptors in the retina (Ehret and Frankenreiter, 1977). These peculiarities are a challenge for the deciphering of the molecular mechanisms underlying the development, maintenance, and function of the auditory organ.

The limited number of cell material as well as the complicated accessibility of the cochlea in the temporal bone have long hampered molecular advances in the auditory field. Recent improvements in the microfluidic sorting of isolated live cells facilitated the application of single-cell RNA sequencing (scRNA-seq) techniques up to thousands of cells and led to new insight into the molecular physiology of cochlear function. Several single-cell transcriptomic studies have investigated the transcriptional diversity of cochlear cell types like in the lateral wall (Korrapati et al., 2019; Gu et al., 2020; Koh et al., 2023), the gene expression profiles of the various supporting cell-types (Hoa et al., 2020), and the heterogeneity and connectivity of both immature and mature auditory neurons (Petitpré et al., 2018, 2022; Shrestha et al., 2018; Sun et al., 2018; Sanders and Kelley, 2022). Lastly, wider scale studies were carried out in developing and young adult cochleae (Kolla et al., 2020; Xu et al., 2022; Jean et al., 2023; Rose et al., 2023), during aging (Sun et al., 2022; Boussaty et al., 2023), and under pathophysiological conditions such as acoustic trauma (Milon et al., 2021). Nevertheless, these extensive transcriptomic characterizations of the cochlea, mostly available to the scientific community (Orvis et al., 2021), are based on dissociated tissues and do not take into account the spatial organization of the cochlea. The cochlear neurosensory epithelium is organized along two major anatomical axes of development, the medio-lateral (radial) axis, that supports differentiation of sensory cells and their surrounding supporting cells (SCs), and the longitudinal (tonotopic) axis that underlies tonotopic properties of the cochlea. The combination of gene expression gradients along these two axes defines unique physiological attributes of the cells intimately linked to their position within the cochlea. A previous work combining microarray and fluorescent in situ hybridization of the transcripts has put in evidence a list of genes differentially expressed between the base and apex of the mouse cochlea on P0 and P8, classifying genes according to their spatial and temporal changes of expression (Son et al., 2012). More recently, a study built a quantitative spatial map of the neonatal organ of Corti based on an expression microarray of 192 genes and known expression patterns of cochlear-specific genes (Waldhaus et al., 2015).

The differential gene expression between the different cochlear cell types and within the same cell type along those radial and longitudinal gradients remains to be characterized. The combination of fluorescent in situ hybridization of the transcripts (FISH) and of scRNA-seq has already allowed the establishment of a list of differentially expressed genes (DEGs) along the tonotopic axis in the tympanic border cells (TBCs). Taking advantage of the tonotopic gradient of Emilin2 (Elastin MicrofibriL INterfacer 2) expression, the ordering of TBCs along the tonotopic axis allowed the identification of other DEGs following similar tonotopic gradients with stronger expression at the base or opposite with stronger expression at the apex (Jean et al., 2023). However, the lack of knowledge of such markers for the other cochlear cell types either along the radial or the longitudinal axes prevents the generalization of this methodology with scRNA-seq. The Xenium in situ plateform (10x Genomics), capable of using up to 5000 probes simultaneously, was recently used to map the desired set of transcripts onto cochlear nuclei-stained sections. This work demonstrated this technique as particularly useful for validation of scRNA-seq experimental results (Hertzano, unpublished). However, the possibility of investigating the locations of thousands of genes simultaneously thanks to high throughput scRNA-seq technologies was not possible until recently.

In this study, we opted for the Visium HD technology (10x Genomics), a sequencing-based technology that employs arrays of spatially barcoded probes to capture RNA molecules unbiasedly from histological tissue sections, achieving transcriptomic mapping at single-cell resolution. We investigated spatial transcriptional stage during embryonic (E) stages, E14 and E16, as well as during postnatal (P) development P1 and P8. Combining these spatial transcriptomics data with classical scRNA-seq (10xGenomics), we not only confirmed the cell-type assignment from the scRNA-seq data, but also refined the classification for some cell-types. Moreover, we put in evidence gradients of gene expression along the tonotopic as well as radial axis in several cell-types together with their temporal dynamics during development.

## Results

### Establishing a single-cell spatially resolved transcriptomic dataset of the mouse cochlea during development

With the aim of optimizing gene detection sensitivity and preserving single-cell resolution, the datasets were analyzed using a binning of 8 µm² based on the 2 µm x 2 µm barcode resolution, on a 6.5 mm x 6.5 mm array, allowing to process up to 5 inner ear sections (each from different mice) simultaneously. Formalin-fixed paraffin-embedded (FFPE) samples stained with hematoxylin and eosin (H&E) were considered to better preserve the inner ear architecture. A total of three inner ear sections on E14, three on E16, four on P1 and four on P8 were analysed, resulting in a total of 21 419, 30 753, 31 176 and 40 599 filtered bins for each age, respectively, were analyzed (see material and methods, Fig. 1, S1, S2, S3, S4). In parallel, we carried out scRNA-seq from E16 onward as HCs have not yet differentiated along the whole cochlear axis on E14 (ref). In parallel, to complement our previously published scRNA-seq atlas on P8 including 26 169 cells (Jean et al., 2023), we carried out additional scRNA-seq experiments on E16 and P1 stages when all hair cells along the cochlear axis have differentiated, resulting in a total of 41 385 and 35 210 cells that passed the quality control filtering, respectively (Fig. 1A, B). For both the Visium HD and scRNA-seq datasets, assignment of cochlear cell-types was carried out based on the analysis of DEGs between the cell clusters. We first looked for expression of canonical marker genes and then validated each assignment by comparing the similarity of the gene expression profiles to published datasets at similar developmental stages available on the umgear database (Kolla et al., 2020; Rose et al., 2023, see materials and methods, Dataset S1). Within the scRNA dataset, the circulating cells (mainly blood cells) were excluded from further analysis, whereas the non-circulating cells that are resident cochlear cells were classified into main ensembles based on their cochlear subregion of origin (Fig. 1B).

**Fig. 1.**
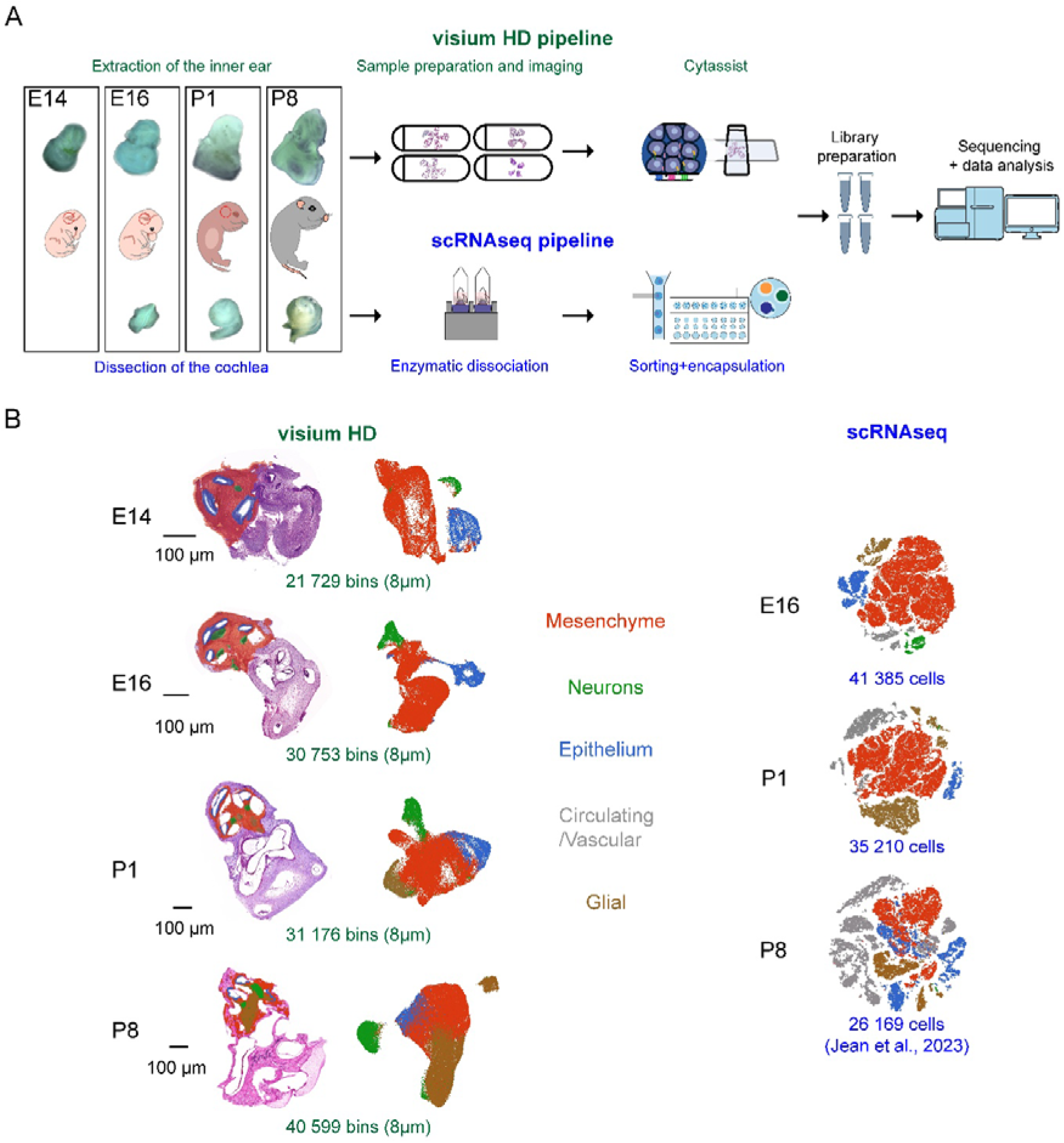
Single-cell transcriptomic characterization of the mouse cochlea during development. (A) Experimental design of the scRNA-seq and Visium HD studies on E14, E16, P1 and P8. (B) Left: T-SNE depicting the scRNA-seq datasets. Right: Visium HD datasets consisting in FFPE histological sections stained with H&E and overlayed with the filtered bins color-coded by the cell-ensemble they were assigned to in the corresponding UMAP plots. No scRNA-seq experiments were performed on E14 whereas the P8 scRNA-seq dataset was taken from Jean et al., 2023.

We first analyzed the embryonic stages E14 and E16. On E14 (Fig. S1), the Visium HD pipeline allowed to distinguish within the epithelium the cochlear roof and the cochlear floor, the latter composed of the greater epithelial ridge (GER) located medially and the lesser epithelial ridge (LER) located laterally. Both the GER and LER contain transient cell populations that regress during post-natal cochlear maturation, leading to the formation of the inner and outer sulcus, respectively, around P8 (Kubota et al., 2021; Kelley, 2022). On E16 (Fig. S2), the cochlear roof could be subdivided into two parts classified as the future Reissner’s membrane and stria vascularis, respectively. The cochlear floor was segmented from its medial to lateral part into interdental cells, GER, HCs, pro-sensory domain and LER (Fig. S2).

The otic mesenchyme cells (OMCs), the most abundant cell-type in the developing cochlea, were detected on E14 and could be subdivided into five cell subtypes on E16 following the nomenclature defined in the study that uncovered their heterogeneity (Rose et al., 2023). Based on their DEGs in the E16 scRNA-seq dataset, we could identify the OMC progenitors and predict the structure they will give rise to including the type I OMCs that will give rise to TBCs secreting the basilar membrane clustered together with the type II OMCs that will develop into the spiral limbus, the type III OMCs that will form the modiolar bones, and the type IV OMCs that will make the fibrocytes of the lateral wall. In the spatial transcriptomic dataset, these mesenchymal cell types were more challenging to distinguish as their clusters merged together (Figure 2B). We could establish that the signature of these clusters was in agreement with their structural location as the type I-IV cells were found surrounding the epithelium, the type II-III cells were detected mainly at the medial part surrounding the types I-IV cells, while the type III cells were filling up the rest of the cochlea. Moreover, on both E14 and E16, a cell-type encapsulating the cochlea named hereafter ‘Shell’ was detected. This cell type was absent in the scRNA-seq dataset because it was removed during the dissection. Finally, the Schwann cells, melanocytes (incorporating the stria vascularis to form the intermediate stria cells (Renauld et al., 2022)), and neurons were detected in the E16 scRNA-seq dataset. As previously reported (Petitpré et al., 2022), we could classify the neurons into two subpopulations based on their fate of projections to the HCs, the Ia/Ib/II and the Ic subgroups. Because the depth of sequencing of the Visium HD data was lower than the scRNA-seq data, these neuronal subgroups could not be detected in this dataset.

**Fig. 2.**
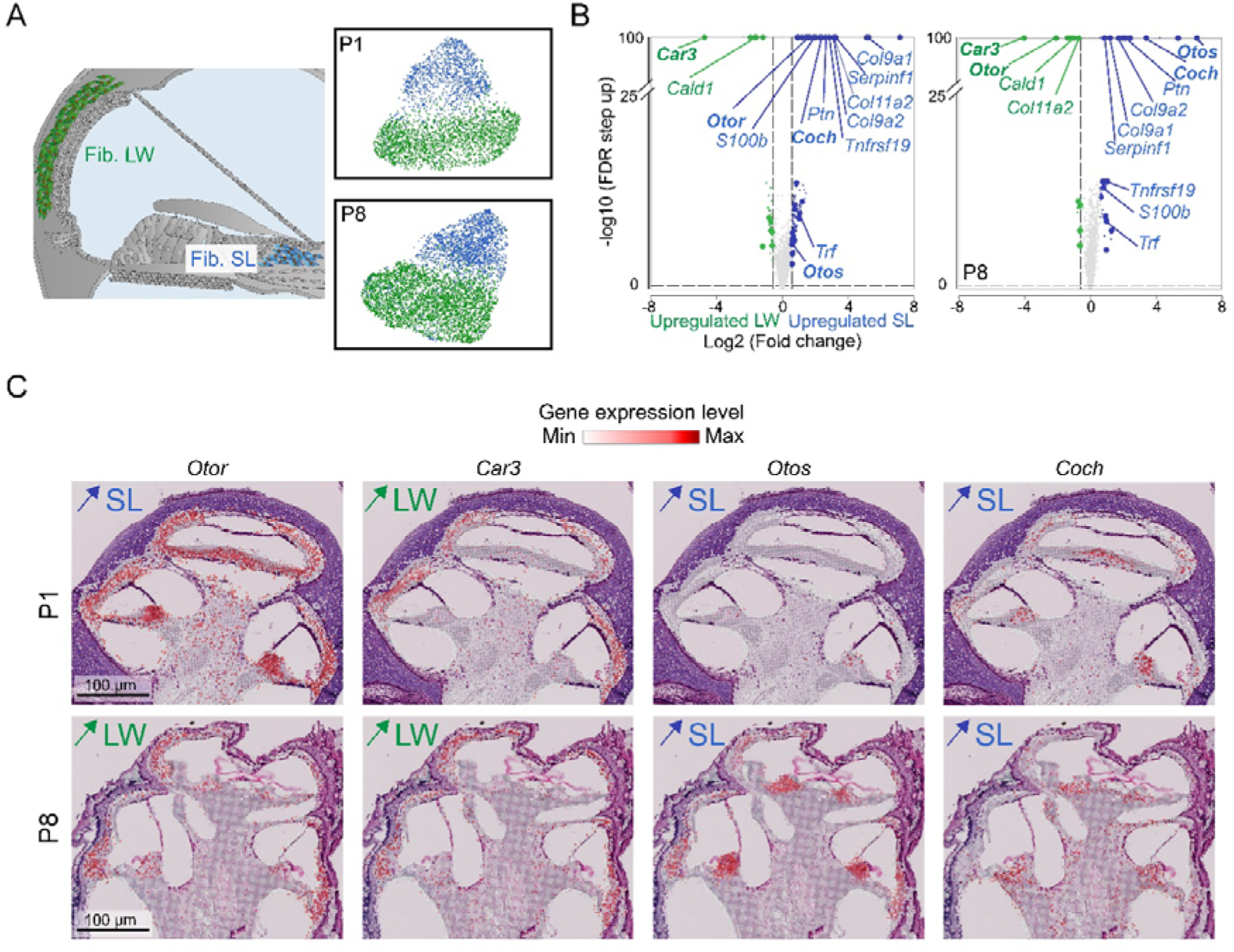
Spatial differential gene expression analysis between fibrocytes of the spiral limbus and of the lateral wall. (A) Left: Diagram illustrating the localization of fibrocytes in the spiral limbus (colored in blue) and in the lateral wall (colored in green). Right: Visium HD UMAP plot of the fibrocyte subtypes on P1 and P8. (B) Volcano plots displaying the expression of all detected genes. Genes with a stronger expression in the spiral limbus are colored in blue, while those with a stronger expression in the lateral wall are labelled in green. The genes belonging to the top 100 DEGs in the fibrocytes whatever the cochlear region are displayed with a bigger spot. The genes highlighted in the manuscript are annotated. The genes with a fold change of gene expression below 1.5 and or a statistical significance above 0.05 are in grey. (C) Histological Visium HD cross sections from FFPE samples stained with H&E, overlayed with the filtered bins color-coded according to the gene expression level (red scale). SL: spiral limbus; LW: lateral wall; Fib: fibrocytes.

We then assigned the cell types analyzed in postnatal cochlear samples. In the P1 scRNA-seq dataset (Fig. S3), the epithelial floor was subdivided from its medial to lateral parts into interdental cells, GER cells, inner phalangeal and border cells, HCs, Deiter’s and pillar cells, and LER cells. These cell-types were also detected in the Visium HD P1 dataset at their expected histological localization, at the exception of the inner phalangeal and border cells that were intermingled within the GER cluster. In the P8 Visium HD dataset (Fig. S4), the GER and LER cells developed into the medial (gathering inner sulcus, phalangeal and border cells) and lateral SCs (assembling the Hensen’s, Claudius’ and outer sulcus cells). The medial part of the epithelial roof that gives rise to the Reissner’s membrane by P1 was detected in the P1 and P8 Visium HD sections. Regarding the lateral part of the epithelial roof that gives rise to the stria vascularis, both marginal and intermediate stria cells were identified in the P1 scRNA-seq data, but not the basal ones. Of note, the basal stria cells were barely detected in our P8-12-20 scRNA-seq dataset, most probably due to bias in the cytometry-based cell sorting and their lack of known markers (Jean et al., 2023). In both the P1 and P8 Visium HD datasets, a cell cluster corresponding to the stria vascularis as well as another one corresponding to the root and spindle cells could be identified. However, their spatial proximity and the limited number of detected genes prevented to separate them further. (Fig S3, 4).

Because the otic mesenchyme has differentiated by P1, the three types of bone cells –the pre-osteoblasts, osteoblasts and osteocytes could be detected in both the scRNA-seq and Visium HD datasets. The lower sequencing depth of the P8 Visium HD dataset, however, prevented the detection of these three cell types, which were classified as “Osseus” (Figure S4). In addition, we had identified in our published P8-P12-P20 scRNA-seq atlas and confirmed through RNAscope assays the presence of two previously undescribed bone-related cell-types in the modiolus we named Surrounding Structures 1 and 2 (SS1 and SS2, respectively) (Jean et al., 2023). However, we had been unable to determine whether those two cell-types represented distinct cell types or different stages of differentiation of the same lineage. In our P1 scRNA-seq dataset, SS2 cells clustered together with osteocytes, whereas SS1 cells were closely related to TBCs. This pattern indicates that SS1 and SS2 do not share a common origin and instead represent two distinct cell-types. SS2 cells also clustered together with the osteocytes in the Visium HD dataset, while SS1 cells were not detected. In both the P1 and P8 Visium HD datasets, TBCs and fibrocytes were also detected and non-assigned mesenchymal cells were simply classified as OMCs. Lastly, the glial cells and neurons were identified in the P1 scRNA-seq dataset. The Schwann cells as well as the proliferating ones were detected, alongside the satellite glial cells that wrap the neuron cell bodies, and their precursors. The neurons were also classified but in a very limited number, preventing the identification of their subtypes.

### Spatial transcriptomics reveals differential gene expression along the medio-lateral axis within several cochlear cell types

Identifying the subpopulations of a given cell-type without spatial cues is a challenge in the transcriptomic field due to their very close signature and lack of differentiating markers. We first investigated whether such spatial cues were present in fibrocytes, which form within the cochlea loose connective tissues, are crucial for maintaining ion homeostasis and blood flow regulation (Furness, 2019), and are present in both the lateral wall and spiral limbus (Fig. 2A). ScRNA-seq did not allow us to discriminate the fibrocytes from the spiral limbus to the ones from the lateral wall, which were grouped in a single cluster of cells (current study and (Jean et al., 2023)). However, based on histology, Visium HD sections allowed to spatially classify the two populations on both P1 and P8. By performing differential gene expression analysis (see material and methods) between fibrocytes of lateral wall and of the spiral limus, we listed a total of 235 genes on P1 (pooling sections of fourteen cochlear ducts originating from four cochleae) and 88 genes on P8 (pooling sections of thirteen cochlear ducts originating from four cochleae) that were differentially expressed between the two regions (Fig. 2B, Dataset S2). When considering the top 100 DEGs in fibrocytes from both regions, 51 and 29 DEGs were shared on P1 and P8, respectively. The majority of these DEGs were more expressed in the spiral limbus (39 out of 51 on P1, 19 out of 29 on P8) than in the lateral wall. Among the 13 DEGs shared between the two stages, 11 followed the same spatial pattern. While some genes had a stronger expression in the spiral limbus such as the fibrocyte specific genes Otos (encoding otospiralin) and Coch (encoding cochlin), whose mutations cause hearing loss (Robertson et al., 1998; Delprat et al., 2002), other genes followed the opposite pattern such as Car3 (encoding the Carbonic Anhydrase 3)and Cald1 (encoding a calmodulin-binding protein). Notably, two genes shifted from a stronger expression in the spiral limbus on P1 to a stronger expression in the lateral wall on P8, Col11a2 (encoding a collagen related protein) and Otor (encoding otoraplin) (Robertson et al., 2000) (Fig. 2B, C). Our data thus demonstrate that cells of the same cell type localized in different regions of the cochlea may be defined by distinct transcriptomic signatures.

Focusing on the neurosensory epithelium on P1, we then investigated the genes differentially expressed in this structure along the radial axis. We performed a hierarchical clustering of the 208 DEGs along the radial axis that resulted in seven clusters distributed between the GER cells, the Deiter’s-pillar cells and the LER cells (Fig 3A, B, Dataset S3). A first cluster of DEGs was enriched in the GER and included one of the differentiating GER marker Crabp1 (Kubota et al., 2021), the deafness gene Gjb2 encoding connexin 26 (ref), Eya1 involved in the branchio-oto-renal syndrome causing hearing loss (Abdelhak et al., 1997) and the planar cell polarity gene Celsr1 (Curtin et al., 2003). A second cluster of DEGs was preferentially expressed in the Deiter’s and pillar cells and included the fibroblast growth factor receptor Fgfr3 known to be specific of these cell-types (Hayashi et al., 2010), Cep41 encoding centrosomal protein 41 and Id4 encoding the inhibitor of DNA binding 4. The third cluster of DEGs was enriched in LER cells, including Bmp4 encoding the bone morphogenic protein 4, Fst encoding follistatin, Otogl encoding otogelin-like, and Emx2 involved in planar cell polarity (Holley et al., 2010). The four other clusters of DEGs were enriched in two of the three regions. The cluster of DEGs in both the GER and LER included Cnmd encoding the cartilage matrix protein chondromodulin, Epiphycan (Epyc) whose deletion induces elevated hearing thresholds (Hanada et al., 2017) and Btg1 known to regulate cell growth and differentiation. Two clusters of DEGs were found in the cells of the GER and Deiter’s-pillar cells. One with a stronger expression in GER cells including Tecta encoding tectorin alpha and Nr2f1, and the other one with a stronger expression in the pro-sensory domain, including the other tectorial membrane complex encoding genes Tectb and Otog (Rau et al., 1999; Avan et al., 2019), as well as the Notch ligand Jag1 (Jag1) required for sensory progenitor development (Kiernan et al., 2006). Finally, the remaining clusters of DEGs had a stronger expression in both LER and pro-sensory domain cells like Uchl1 that was shown to be downregulated in age-related hearing loss (Li et al., 2023), and Syne2. These DEGs specific of the neurosensory epithelium had similar patterns of expression in the P1 scRNA-seq data (Fig. 3A).

**Fig. 3.**
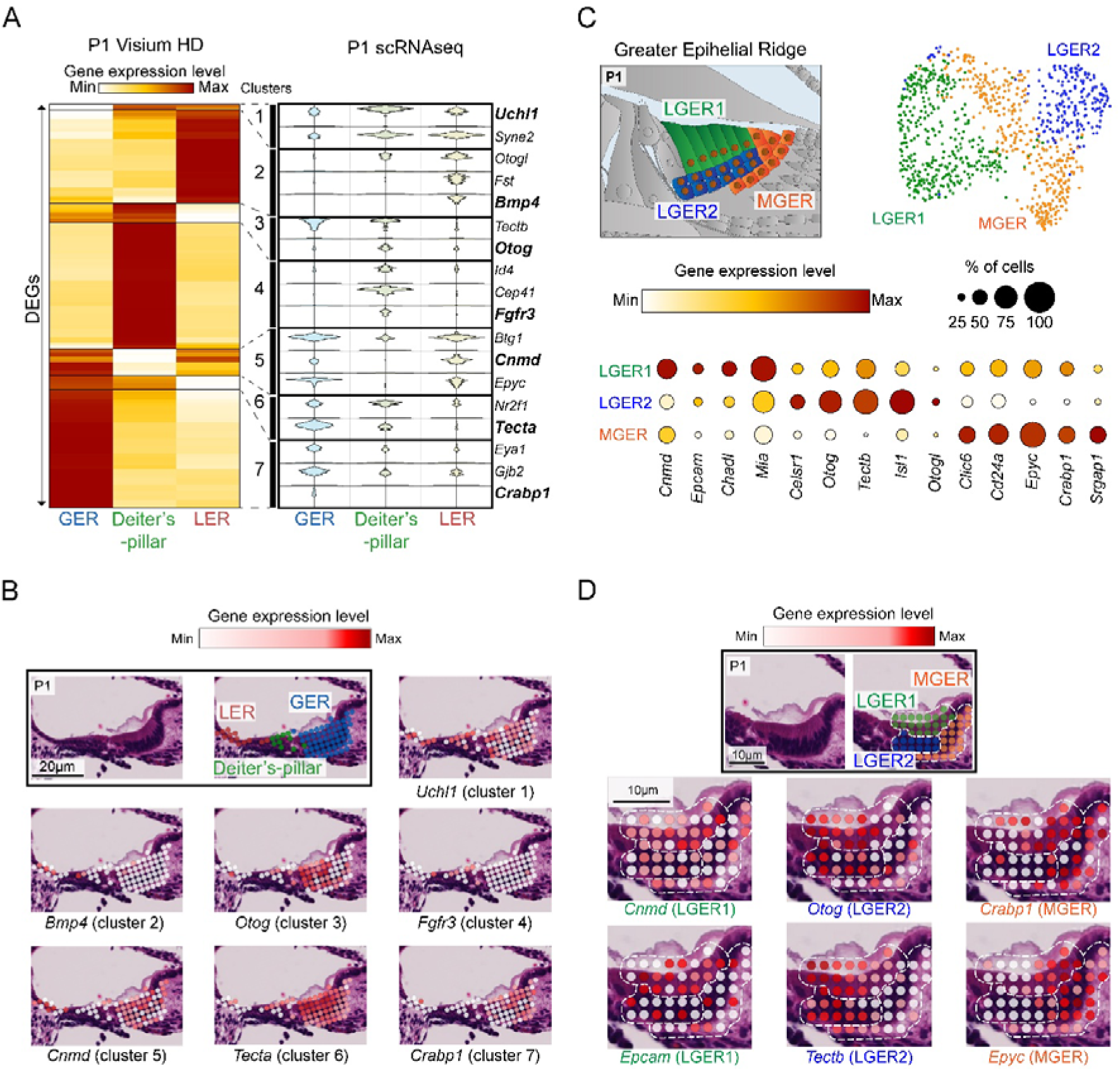
Radial differential gene expression analysis between supporting cell types on P1. (A) Left: Heat map/hierarchical clustering of the DEGs distributed between GER cells, Deiter’s-pillar cells, and LER cells on P1. Right: Violin plots depicting the expression of the DEGs in the Visium HD and scRNA-seq datasets on P1. (B) Histological Visium HD cross sections from P1 FFPE samples stained with H&E, overlayed with the filtered bins color-coded according to the gene expression level. (C) Top left: diagram illustrating a cross section of the cochlea where the cell-subtypes from the MGER, LGER1 and LGER2 are color-coded. Top right: Visium HD UMAP plot of the GER subtypes on P1. Bottom: Bubble plot analysis comparing the expression levels of a subset of DEGs between the three cell subtypes. Gene expression level is color-coded while the proportion of cells expressing the genes is indicated by the size of the spots. (D) Histological Visium HD cross sections from FFPE samples stained with H&E, overlayed with the filtered bins color-coded according to the gene expression level, and delimited by dashed lines depending on the GER cell subtype they belong to. L. SCs: Lateral supporting cells, M. SCs: Medial supporting cells, MGER: Medial greater epithelial ridge. LGER1 & 2: Lateral greater epithelial ridge 1 & 2.

A similar analysis was carried out on our P8 visium HD dataset. Six clusters of DEGs (90 genes in total) were distributed between the medial SCs, the Deiter’s-pillar cells and the lateral SCs (Fig. S5). A first cluster of DEGs with a stronger expression in the lateral SCs included the bone morphogenic protein encoding genes Bmp4 and Bmp6, and the cytoprotective clusterin gene Clu (Lee et al., 2017). The Deiter’s-pillar cells differentially expressed Cep41, Fgfr3 and Id4, similarly like in the P1 data set, as well as the tectorial membrane related genes Otog and Otogl. The medial SCs differentially expressed Clic6 encoding a chloride intracellular channel and Mia encoding the melanoma inhibitory activity protein. The remaining clusters had DEGs expressed in two cell-types as for the medially and laterally located SCs that differentially expressed Epyc, Cnmd, collagen related genes (Col9a1/2/3, Col11a2) and Lum encoding Lumican, a regulatory proteoglycan of collagen-rich tissues, or as for the pro-sensory domain and lateral SCs that differentially expressed kcnj16 encoding the potassium channel kir5.1 and Fbxo2 whose deletion induces accelerated age-related hearing loss (Nelson et al., 2007). Finally, the expression of the Tecta and Tectb genes, as well as Ush1c encoding the mechanoelectrical transduction component harmonin were enriched in the medial SCs and pro-sensory domain in comparison with the lateral SCs. In our previously published P8 scRNA-seq dataset (Jean et al., 2023), the SCs from the medial and lateral parts could not be separated, we are now able to differentiate them with the markers found in the Visium HD dataset (Fig. S5).

We then investigated transcriptomic differences within subregions of the GER on P1 (Fig 3C, D). Previous non-spatially resolved scRNA-seq studies put in evidence in mice and rats different cell subtypes within the GER along its radial axis taking advantage of in situ hybridization assays and already published positional databases (Kolla et al., 2020; Chen et al., 2021a). In our dataset, three cell subtypes of the GER could be isolated, one in the medial region of the GER (MGER), and two others in the lateral regions of the GER (LGER1 and 2), with LGER2 located underneath LGER1. Of note, their relative localization to each other along the radial axis was identical in all the analyzed cochlear epithelia along the tonotopic axis. By analyzing the top 100 DEGs in the GER, we extracted 31 markers differing the most between the three GER cell subtypes. DEGs in the MGER included Crabp1 (Kubota et al., 2021) and Epiphycan (Epyc). LGER1 cells differentially expressed epithelial cell markers such as Cnmd encoding Chondromodulin (Jan et al., 2021) and Epcam, whereas LGER2 cells located underneath expressed DEGs encoding proteins of the tectorial membrane including Otog (encoding otogelin), Otogl encoding otogelin-like and Tectb (Avan et al., 2019; Gagliardini et al., 2025).

### Spatial transcriptomics resolves tonotopic gene expression gradients in several cochlear cell-types along the longitudinal axis

In our published P8 scRNA-seq dataset obtained from P8 mice (Jean et al., 2023), we had taken advantage of the increasing tonotopic gradient of Emilin2 expression from the apex to the base of the cochlea to split TBCs into four equal-sized groups of cells. According to Emilin2 expression levels, TBCs were had been assigned to the base (B), middle-base (MB), middle-apex (MA), and apex (A) of the cochlea in the P8 scRNA-seq dataset. This analysis established a list of DEGs displaying either a base to apex gradient of expression (strongest at the base, B>MB>MA>A), or an apex to base gradient of expression (strongest at the apex, A>MA>MB>B), some of which were confirmed by RNAscope assays (Jean et al., 2023).

Regarding the Visium HD datasets, the cochleae were also subdivided into four tonotopic regions based on histology, thereby roughly mirroring our previous nomenclature, the DEGs differing the most between the apical and basal cells of all analyzed cochleae were included (see material and methods). For a given cell type, only the top 100 DEGs were considered. On P8, the tonotopic analysis was based on the differential analysis between the middle-apex and base of four cochleae, among which only one showed an apical part displayed in the data (Fig. 4A)we isolated two other cell-types (Fig. S6, Dataset S4) with DEGs showing not only a significantly stronger expression either at the base or at the apex, but also a strict tonotopic gradient meaning that their mean expression level decreased or increased gradually from base to apex (MA>MB>B or B>MB>MA). Concomitantly with increasing gene expression levels, the proportion of bins expressing the gene also increased in most cases. Regarding TBCs in the P8 Visium HD dataset, genes with a strict base to middle-apex gradient (B>MB>MA) that were already identified by scRNA-seq (Jean et al., 2023) like Emilin2 and Rarres1 were validated. However, the down-regulator of Wnt signaling Notum, showed a gradient of expression (MA>MB>B) opposite to the one observed in our P8 scRNA-seq data (Jean et al., 2023) (Fig. 6B). In the SGNs, 16 DEGs were identified, all presenting a base to middle-apex gradient of expression (B>MB>MA), including Nefl encoding the neurofilament light chain (Nefl), Calb2 encoding the calcium binding protein calbindin 2, Syt2 encoding the synaptic vesicle membrane protein synaptotagmin 2and Eef1a2 encoding a translation elongation factor. Finally, HCs and the Deiter’s-pillar cells were pooled together for analysis owing tothe limited number of HCs and bins overlapping both cell-types. Four DEGs followed a tonotopic pattern of expression, two with a higher expression at the middle-apex, the immunoglobulin related gene Lghm, found to be differentially expressed in OHCs (Xu et al., 2022) and Tectb. Conversely, Tmprss33 encoding the transmembrane serine protease 3, expressed in HCs and critical for their survival at hearing onset (Fasquelle et al., 2011), as well as Sh2d6 were more expressed at the base than at the middle-apex.

**Fig. 4.**
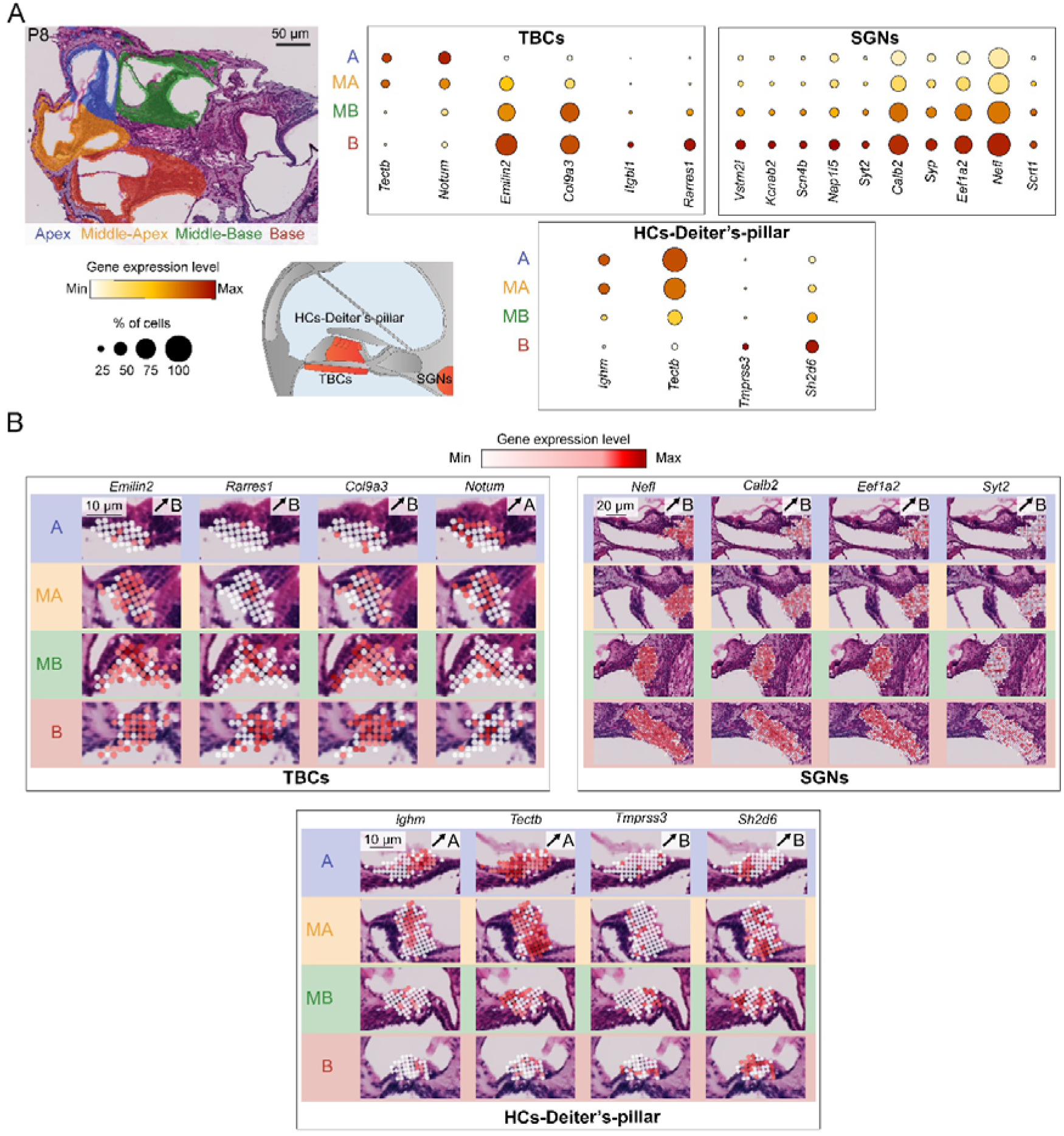
Tonotopic gradient of gene expression in the TBCs, SGNs and HCs-Deiter’s-pillar cells on P8. (A) Top left: Histological Visium HD cross sections from FFPE samples stained with H&E, showing a cochlea analyzed, overlayed with the filtered bins color-coded according to their tonotopic location being the apex in blue, middle-apex in yellow, middle-base in green and the base in red. Bottom left: Cross section schematic of the four cell-types analyzed. Right: Bubble plot analysis comparing the expression levels of a subset of genes with a tonotopic gradient of expression either from base to apex, or apex to base. The gene expression level is color-coded, while the proportion of cells expressing the genes is indicated by the size of the spots. (B) Histological Visium HD cross sections from FFPE samples stained with H&E, overlayed with the filtered bins color-coded according to the gene expression, at the different tonotopic localization for each of the analyzed cell-types.

**Fig. 5.**
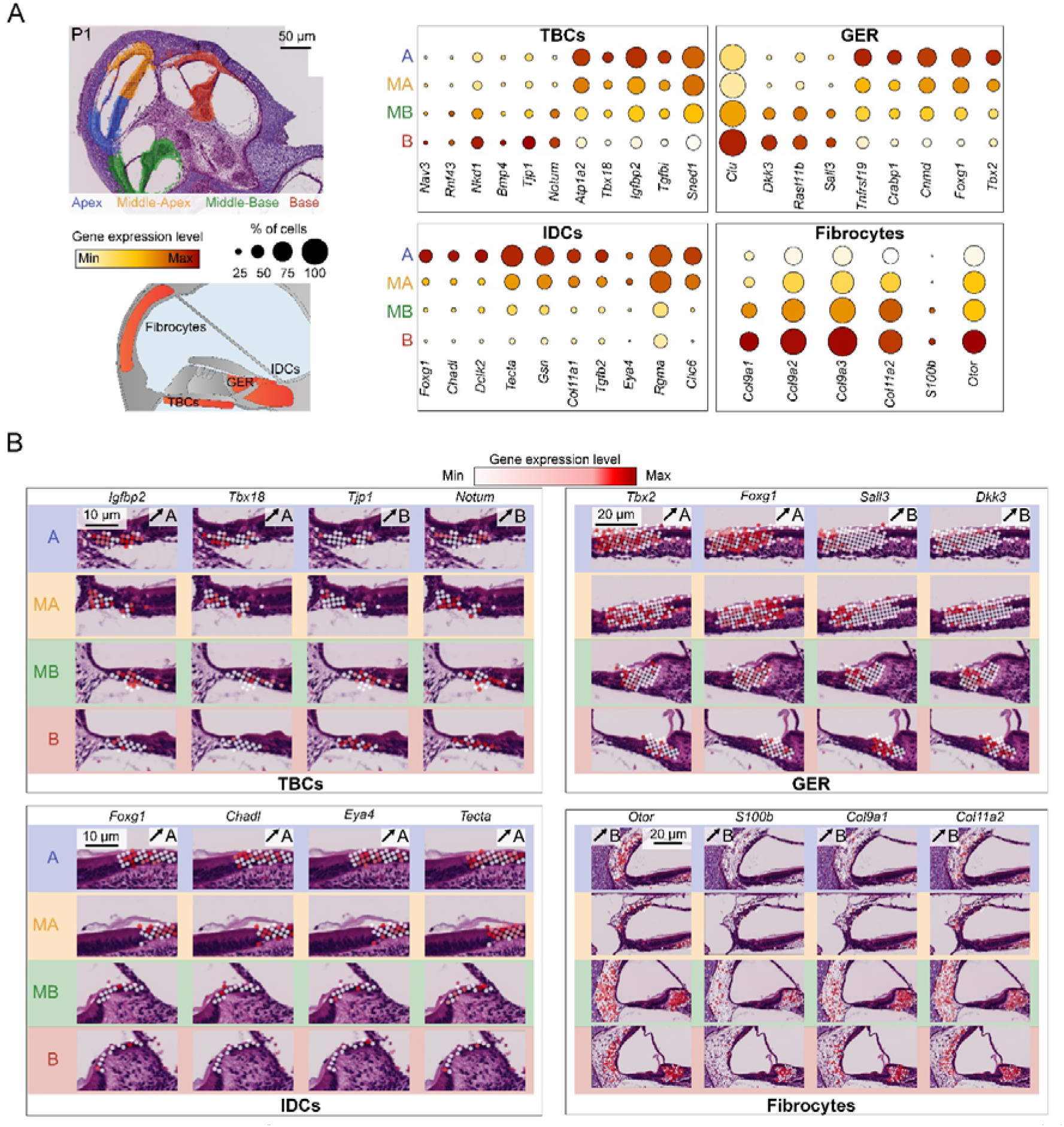
Tonotopic gradient of gene expression in the TBCs, GER cells, interdental cells and Fibrocytes on P1. (A) Top left: Histological Visium HD cross sections from FFPE samples stained with H&E, showing a cochlear tissue analyzed, overlayed with the filtered bins color-coded according to their tonotopic location at the apex (in blue), middle-apex (in yellow), middle-base (in green) and base (in red). Bottom left: Diagram illustrating the four cochlear cell-types analyzed. Right: Bubble plot analysis comparing the expression level of a subset of genes having a tonotopic gradient of expression either from base to apex, or from apex to base. The gene expression level is color-coded while the size of the spots indicates the proportion of cells expressing the genes. (B) Histological Visium HD cross sections from FFPE samples stained with H&E, overlayed with the filtered bins color-coded according to the gene expression, at the different tonotopic localization for each of the cell-types analyzed.

**Fig. 6.**
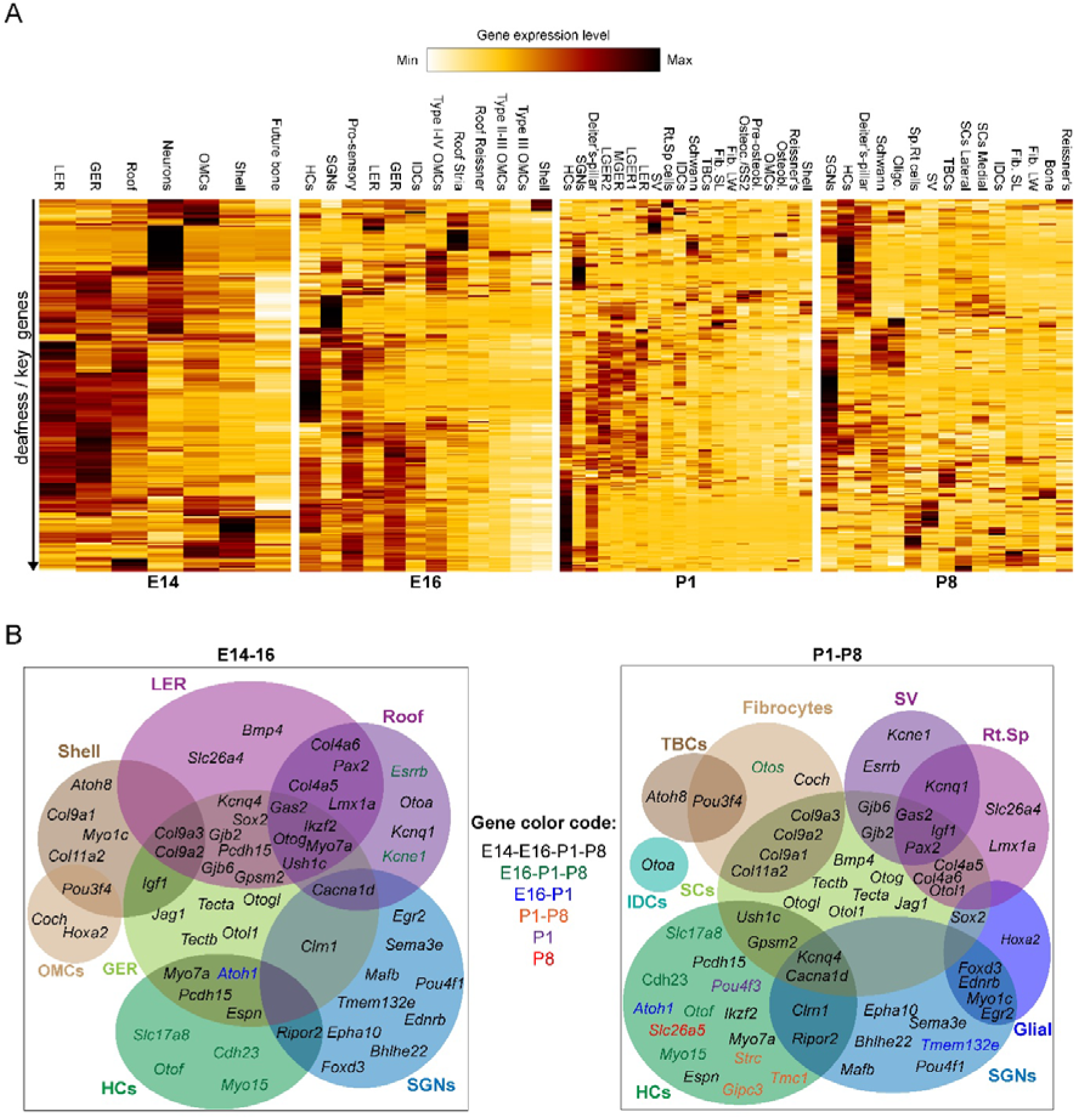
Cochlear cell expression pattern of deafness and key genes along development. (A) Heat map/hierarchical clustering of all detected deafness and key genes and cell types at all stages. (B) Distribution of deafness genes and key genes between the cell types at embryonic (left) and post-natal stages (right). Genes are color coded according to their stages of expression.

On E16, the tonotopic analysis was performed on the one cochlea that presented the four tonotopic parts (Fig. S7). Two other cochleae only showed the apical and middle-apical parts, and were excluded from the analysis. However, the tonotopic gradients of gene expression identified in the complete cochlea were observable in those two excluded cochleae (Fig S8). We isolated four cell-types that presented a list of strict tonotopic DEGs.) (Fig. S9, Dataset S4). 14 DEGs were isolated for GER cells. Among them, 4 genes were more expressed at the cochlear apex like Epcam, a marker of epithelial cells, and the planar cell polarity gene Celsr1 (Curtin et al., 2003). In contrast, 10 DEGs had an inverted gradient of expression with a stronger expression at the cochlear base like the deafness gene Myh14 (Kim et al., 2017), the thyroid hormone receptor β Thrb and the differentiating GER marker Crabp1. For LER cells, 21 DEGs were found, all of them (at the exception of Galnt7) being more expressed at the apex like the inhibitor of DNA-binding 2 Id2, the Zinc-finger transcription factor Gata2, as well as the deafness genes encoding the cytoskeletal regulatory protein Gas2 (Chen et al., 2021b) and Sprouty 2 (Spry2) (Shim et al., 2005). By combining the cells from the epithelial roof (both Reissner membrane’s and stria regions), 10 DEGs were uncovered, 3 of them displaying a stronger expression at the apex (Oc90, Sfrp1 and Ptprf), while the 7 others were more expressed at the base like Gas2 (showing an inverted gradient as compared to LER cells), as well as the doublecortin-like kinase 1 Dclk1 and Enpep, both markers of marginal stria cells at later stages. Finally, a total of 23 DEGs were identified in the SGNs, 3 of them with a stronger expression at the base (Lrrc55, Stac2 and Snhg11), while the other genes were more expressed at the apex (Shox2 and Nrg3). No DEGs in interdental cells and HCs-pro-sensory domains were found when comparing cochlear apical and basal regions (Fig. S9).

We could detect the largest number of cell types with tonotopic gradient on P1, six of them. (Fig. S10, Dataset S4). Similarly to E16 and P8, only the genes presenting a strict tonotopic gradient with a stronger expression at the apex were considered (apex to base gradient: A>MA>MB>B). For opposite gradient, the filtering was less strict and included the genes showing either a B>MB>MA>A or B>MB>A>MA pattern of expression. Unlike on E16, DEGs with a strict base to apex gradient of expression (B>MB>MA>A) were rarer, most of them having a slightly higher expression at the apex than at the middle-apex of the cochlea. In the TBCs, 18 DEGs with a strict apex to base gradient were found including the genes encoding the ATP pump subunit (Atp1a2), the T-Box transcription factor (Tbx18), the insulin growth factor binding protein (Igfbp2) and the extracellular matrix Insulin Responsive Sequence DNA Binding Protein-1 (Sned1). Regarding the DEGs with the opposite strict gradient (B>MB>MA>A), we detected the genes encoding the neural morphogenesis protein Nav3, as well as Notum, Nkd1 and Rnf43, all three of which encode negative regulators of Wnt signaling (Kakugawa et al., 2015; Larraguibel et al., 2015; Tsukiyama et al., 2015). 5 other DEGs had a less strict base to apex gradient including the ones encoding the bone morphogenic protein 4 (Bmp4) and the tight-junction protein ZO-1 (Tjp1). In contrast to the P8 scRNA-seq data, Emilin2 is weakly expressed on P1 and cannot be used as a proxy for tonotopy. Instead, we took advantage of the P1 Visium HD dataset in which Igfbp2 was detected as a tonotopic marker with a stronger expression at the apex than at the base. This gene was expressed in 92% of the TBCs in the scRNA-seq data (3264 out of 3557 cells), allowing the separation of TBCs into four equal groups, each corresponding to a tonotopic part of the cochlea. The tonotopic DEGs in the P1 scRNA-seq data mirrored most of those in the P1 Visium HD dataset., including genes with an apex to base gradient (Igfbp2, Atp1a2, Tbx18, Tgfbi, Aldh1a2 and Sned1) and genes with a reverse gradient (Notum, Nav3, Nkd1, Tjp1, Akap12), at the exception of Rnf43) (Fig. S11). Several of these DEGs in TBCs of the P1 Visium HD dataset maintain their tonotopic gradient of expression later in development in our published P8 scRNA-seq dataset (Jean et al., 2023) like Notum that has a base to apex gradient, and Sox11, Robo1, Tead2 and Tbx18 that have an apex to base gradient of expression.

We then focused on the GER and identified 17 DEGs with a strict apex to base gradient (A>MA>MB>B) including Foxg1 encoding a transcriptional regulator indispensable for inner ear development (He et al., 2019), Cnmd encoding an epithelial cell marker, Crabp1, and Tbx2 encoding a differentiation factor of HCs and SCs (Kaiser et al., 2022). No DEGs with a strict base to apex gradient (B>MB>MA>A) were detected. However, several genes with a less strict tonotopic gradient were identified including Clu encoding the cytoprotective Clusterin (Lee et al., 2017), Sall3 encoding a zinc finger related protein and Dkk3 encoding a modulator of Wnt signaling. Regarding the interdental cells, a cell type that showed no tonotopic gradient on E16, 17 DEGs along the tonotopic axis were identified on P1. All of them had a higher expression at the apex than at the base of the cochlea. These DEGs along the tonotopic axis included genes encoding the transcriptional regulator Forkhead Box G1 (Foxg1), the collagen binding protein Chondroadherin-like (Chadl), as well as the deafness gene Tecta. When pooling fibrocytes the spiral limbus and lateral wall, we could identify 6 DEGs along the tonotopic axis, all with a stronger expression at the base than at the apex of the cochlea, including collagen related genes (Col9a1, Col9a2, Col9a3, Col11a2) as well as the cochlear genes Otor encoding Otoraplin (Robertson et al., 2000) and S100b encoding the EF-hand calcium-binding related protein. Notably, we observed a similar set of genes on P8 including Col9a1, Col9a2, Col11a2, and S100b. While S100b conserved its strict pattern of tonotopic expression, the strict gradient of expression for collagen related genes was lost although their expression was stronger in basal regions of the cochlea than apical ones (Fig. S12). Pooling together HCs and the Deiter’s-pillar cells for analysis revealed 6 DEGs along the tonotopic axis (Fig. S13). Three DEGs had a strict tonotopic gradient of expression: Fgfr3 encoding the fibroblast growth factor receptor 3 known to be specific of pillar and Deiter’s cells (Hayashi et al., 2010), Ccer2 shown to be HC specific (Kolla et al., 2020), and Tectb. The three other genes had a reverse gradient from base to apex: Calb1 encoding for the calcium binding protein 1 present in HCs, GER and SGNs (Sanders and Kelley, 2022), Pcp4 expressed in HCs (Xu et al., 2022) and Fclrb. DEGs were also detected in SGNs analyzed only from the base to the middle-apex due to the absence of analyzable apical spiral ganglia. 25 DEGs were identified, 7 of them with a stronger expression at the middle-apex including the transcription factor and SGN subtype marker Pou4f1 in agreement with past immunostaining studies (Sherrill et al., 2019), and Celf3. The DEGs with a higher expression at the base than the apex of the cochlea included Nefh encoding the neurofilament heavy chain protein and Calb2 encoding the calcium binding protein calbindin 2/calretinin (Fig. S13).

### A spatial atlas of deafness gene expression along cochlear development

Finally, we investigated along development the spatial clustering of genes involved in deafness or key to cochlear development and function (Elliott et al., 2021, hereditaryhearingloss.org). We analysed the cochlear expression patterns of about 200 detected genes on E14, E16, P1 and P8 (Fig. 6, Dataset S5) that clustered up to ∼20 ensembles on P8. These genes were either specific of a given cell type, expressed in several cell-types, or ubiquitous across development. They could also be expressed in the same cell-type(s), or change their pattern of expression due to the rearrangement of cochlear architecture for instance. Notably, we identified a cluster of genes expressed in the GER at prenatal stages and SCs at postnatal stages, including the ones encoding tectorin-α and tectorin-β (Tecta (DFNA8), Tectb), non-collagenous tectorial membrane proteins (Otol1) and proteins involved in the attachment of the OHC hair bundle to this membrane, such as otogelin (Otog) (DFNB18B) and otogelin-like (Otogl) (DFNB84B). We also detected several genes expressed in the cochlear roof embryonically, that are specific postnatally of the root and spindle cells like Lmx1a (DFNA7) and Slc26a4 encoding pendrin (DFNB4), as well as of the marginal stria cells like the ones encoding the Kcne1/Kcnq1 complex and Esrrb encoding the estrogen-related receptor β (DFNB35). Of note, Gjb2 and Gjb6, known to encode the connexin 26 and 32 (DFN1A and DFN1B), respectively, were found to be expressed in the GER and LER at early stages, and in the SCs and stria vascularis on P8. The fibrocytes co-expressed Pou3F4 (DFNXB) with the TBCs, collagen related genes such as Col9a1,2 and 3 with the SCs, or expressed specifically Otos and Coch (DFNA9). SGNs expressed specifically and robustly several genes at all stages of genes including Pou4F1, Bhlhe22 and Sema3e. In contrast, while HCs expressed the transcriptional regulators of HC differentiation Atoh1 (DFNA89) and Pou4f3 (DFNA15) until P1, these genes were not detected on P8. Conversely, Slc26a5 encoding prestin (DFNB61) and mediating OHC motility (Liberman et al., 2002) was detected on P8 but not at earlier stages. Finally, few genes had the peculiarity to have a broad expression through development, such as Cacna1d encoding a Ca^2+^ voltage-gated channel subunit expressed in cochlear roof, GER and SGNs embryonically, to be then expressed in the SCs, HCs and SGNs postnatally. Similarly, Gas2 (DFNB125), encoding the growth arrest-specific protein 2, was found embryonically in the cochlear roof, LER and GER; to be then expressed postnatally in the stria vascularis, spindle-root cells and SCs. Altogether, this spatial single-cell transcriptomic atlas provides a positional database along cochlear development of the genes indispensable for proper hearing function, a key resource in the proper design of gene therapeutic strategies.

## DISCUSSION

Taking advantage of the Visium HD technology, we established a spatial transcriptomic atlas of the developping mouse cochlea at the single-cell resolution in order to not only validate the results obtained from classical scRNA-seq, but also to gain insights into the differentially expressed genes along the two major axes of development of the cochlea. Unlike scRNA-seq experiments where the cochleae were dissected out, and pooled for subsequent tissue dissociation and cytometry-based cell sorting, the Visium HD pipeline allowed to simultaneously capture every portion of the inner ear without introducing any bias for their transcriptional comparison. We obtained a direct view of the location of the different cochlear cell-types, making it possible their differential gene expression analysis along the radial and tonotopic axes of the cochlea.

The P1 and P8 Visium HD datasets brought new insight into the DEGs along the medio-lateral axis. Focusing first on fibrocytes, we could segregate them into two subtypes based on their localization, the ones from the spiral limbus facing the spiral ganglion, and the ones from the spiral ligament located in the lateral wall. At both developmental stages, we extracted a set of common DEGs more expressed either laterally or medially. Notably, most of these genes conserved their asymmetry of expression with aging. It was known that the fibrocytes were present in those two regions, but there was no transcriptomic evidence allowing their discrimination. We then examined the DEGs among SCs depending on their localization in the neurosensory epithelium, and could hierarchically cluster them. We found seven and six clusters of DEGs, on P1 and P8, respectively, and extracted genes enriched in either one or several of those supporting cell types. The medial and lateral SC markers from the P8 Visium HD data allowed us to refine our previously published P8 scRNA-seq dataset (Jean et al., 2023), where the SCs from the medial and lateral parts could not be segregated apart, and are now distinguishable. Moreover, based on the analysis of the DEGs in GER cells on P1, spatial transciptomics allowed us to divide this transient structure into three subregions, one medial called MGER and two lateral ones superimposed and designated here as LGER1 and LGER2. Remarkably, their relative localization to each other was perfectly maintained in the neurosensory epithelium from the different tonotopic positions and cochleae analyzed.

The Visium HD technique also identified DEGs along the tonotopic axis. Considering only the cell types that could be assigned to a tonotopic position on E16, P1 and P8, thus excluding the mesenchymal and bony cells, we extracted sets of DEGs and observed whether they presented a tonotopic gradient of gene expression level either from base to apex (B>MB>MA>A) or apex to base (A>MA>MB>B). In parallel, the results from Visium HD refined the tonotopic analysis of our P1 scRNA-seq dataset. Considering the wide expression (92% of the cells) of Igfbp2 in the TBCs from the P1 spatial transcriptomics data, and its strict tonotopic gradient of expression intensity from apex to base, we could use this marker as a proxy to divide the TBCs of the P1 scRNA-seq data into the four tonotopic subregions. This approach reinforced the results from the spatial data as the vast majority of specific tonotopic genes in TBCs from the P1 Visium HD dataset followed the same gradient of expression as in the scRNAseq dataset. However, one weakness of this approach is TBCs were assumed to be equally numbered between the four tonotopic positions, which is inaccurate but hardly quantifiable. Regarding the P8 Visium HD datasets, several DEGs were found in common with the P1 Visium HD and scRNA-seq data sets. Nevertheless, the lower number of reads per bin obtained on P8, probably due to increased tissue fragility, prevented to carry out an analysis as deep as for the earlier stages.

Finally, considering only the genes involved in congenital deafness as well as in development and function of the cochlea, we were able to establish a spatial atlas of their expression across development. We identified sets of genes either conserving their pattern at all stages as the ones specifically expressed in SCs (Otog, Otogl, Otol1, Tecta, Tectb, Jag1) and SGNs (Pou4f1, Bhlhe22, Sema3e, Mafb, Epha10), or refining their expressions with the development of cochlear architecture. For instance, genes enriched in the cochlear roof (Lmx1a, Kcnq1, Kcne1, Essrb) were then found to be specific either of the stria vascularis cells (Kcne1, Esrrb), root and spindle cells (Lmx1a), or both (Kcnq1).. Other genes were conservely expressed in several cell-types across development, such as Cacna1d (HCs-SCs-SGNs). As expected in HCs, we confirmed that developmental genes such as regulators of HC differentiation such as Atoh1 and Pou4f3 dissapeared postnatally, or conversely as for the motor protein prestin (encoded by Slc26a5), that was only detected on P8.

Single-cell spatial transcriptomic techniques such as Visium HD undergo non-negligeable limitations due to the intrinsic characteristics of the tissues. In this study, we opted for 5 µm-thick sections, assuming that a single spatial barcode will capture the transcriptome of a single cell in the Z axis. However the cell body placements and thicknesses are variable within any tissue, therefore the spatial barcode might capture the RNA of two adjacent cells or even two different cell types on top of each other. However, since the cochlea is a structure organized helicoidally where cell-types are systematiccaly arranged on the longitudinal axis, two different cell-types are unlikely to be simultaneously captured on a transversal tissue section. Additionaly, Visium HD suffer from its lower capture of mRNA transcripts as compared to scRNA-seq, resulting in a loss of information for genes with relatively low expression levels. Moreover, their lower transcriptomic coverage makes it harder to distinguish similar cell-types. In our study, the binning of 8 µm² (instead of 2 or 16) was the best compromise to jungle between a single-cell resolution and a high enough quantity of captured RNA to assign the bin to a cell-type and perform differential gene expression analysis. However, these bins can be bigger than small cell-types, and delineating the cochlear cells in H&E sections is challenging since only the nuclei and cytoplasms are stained, and not the membranes. Adding to the fact that the cell size and morphology are highly variable in the cochlea, it is impossible to assign every bin to their corresponding cell type without overlaping to one another, compromising the resolution and accuracy of differential gene expression. The perfect example is the stria vascularis, where the respective markers of the basal, intermediate and marginal cell layers could be identified, but could not be assigned to specific bins. Additionaly, we were unable to distinguish the spindle and root cells, the inner phalangeal/inner border cells and GER cells that are transcriptomically and spatially closely related.

In conclusion, despite their lower depth of sequencing that will be improved in the coming years, these spatial transcriptomic datasets along development are an essential contribution to the classic scRNAseq studies due to their direct and unbiased view of the transcripts localization on the tissues. The insights brought by this technique regarding the medio-lateral and tonotopic gradient of gene expression are crucial to fully comprehend the development, maintenance and function of the auditory organ. Moreover, understanding the temporal and spatial gene expression at the single-cell level in the cochlea is a key step for the optimization of therapeutic procedures to ensure efficient and safe therapies with durable effects in order to restore cochlear structure and function.

## ACKNOWLEDGMENTS

This work was supported by grants from the EMBO (ALTF 852–2019 long-term fellowship awarded to P.J., the Institut Pasteur for the Pasteur Roux Cantarini postdoctoral fellowship to S.M., the Agence Nationale de la Recherche “AUDINNOVE” (ANR-18-RHUS-0007 to C.P.),“ Laboratoire d’Excellence “LIFESENSES” (ANR-10-LABX-65), the “NEUROPATHEAR “ (ANR 24 CE16 4488 01 to N.M.), the “SelfMorphogEar” (ANR-21-CE13-0038-013-01 to R.E.), “the France 2030 program” (ANR-23-IAHU-0003), the Raimonde and Guy Strittmatter Foundation, the CARNOT initiatives, the Institut Pasteur (PTR 618–23 to N.M. and M.H.), and the “Fondation pour l’Audition”(FPA IDA05 to C.P. and FPA IDA03 to N.M.). We acknowledge the support of the Fondation pour l’Audition to the Institut de l’Audition. We also would like to thank the Biomics plateform from Institut Pasteur for the sequencing of the libraries, and Sensorion’s contribution for the purchase of the ScRNA-seq kits.

## Author contributions

P.J., S.M. and N.M. designed research; P.J., S.M., A.S-E., A.G., L.B-C., A.L., S.Megharba., performed research; P.J. and C.T. analyzed data; S.N., C.M-C., D.H., A.M., M.H., R.E., C.P., and N.M. supervision; P.J. and N.M. wrote the paper.

## MATERIALS AND METHODS

### Animals

C57BL6/J wild-type mice were purchased from Janvier Laboratories. Animal experiments were performed in accordance with French and European regulations for the care and protection of laboratory animals (EC Directive 2010/63, French Law 2013–118, February 6, 2013), under authorizations from the Institut Pasteur’s ethics committee for animal experimentation.

### Tissue preparation and sample sequencing processing

For scRNA-seq, the cochleae of E16 and P1 animals were extracted by removing the osseous labyrinth and dissected in phosphate-buffered saline (PBS) solution. Briefly, after removal of the osseous shell, whole cochleae were microdissected and then subjected to enzymatic dissociation (Adult Brain Dissociation kit, 130-107-677, Miltenyi Biotec) and mechanical trituration with a gentleMACS dissociator (Miltenyi Biotec). The resulting suspension was filtered sequentially through 70 µm and 40 µm meshes and then sorted by flow cytometry on the basis of size (forward scatter), granularity (size scatter), and viability with a BD FACSAria III cell sorter (BD Biosciences) in PBS with 0.04% bovine serum albumin (BSA). The sorted single-cell suspension was treated (approximatively 3 h after animal sacrifice) according to the “Chromium Next GEM Single Cell 3’ v3.1 protocol” (CG000204 Rev D, 10x genomics) in accordance with the manufacturer’s instructions, for library construction.

For Visium HD, the cochleae of E14, E16, P1 and P8 animals were extracted in phosphate-buffered saline (PBS) solution, and fixed by immersion in 10% Formalin in PBS for 1H at room temperature. Only the organs on P8 were decalcified by incubation in 0.35 M EDTA (ethylenediaminetetra-acetic acid) in PBS (pH 7.5) for 24H at 4 °C. Tissues were then washed in PBS, and dehydrated in ethanol (70% EthOH 1H at 37°C, 70% EthOH 1H at 37°C, 100% EthOH 30 min at 37°C, 100% EthOH 30 min at 37°C, 100% EthOH 1H at 37°C), to be then cleared in isopropanol at 40°C for 1H and embedded in paraffin at 62°C for 4-5H. Five-μm thick paraffin sections were placed on superfrost plus slides and stained with Haematoxylin and Eosin (H&E). Slides were scanned using the AxioScan 7 (Zeiss) system and images were analysed with the Zen 2.6 software. The samples were then processed, in accordance with the manufacturer’s instructions, using the “Visium HD FFPE Tissue Preparation Handbook, CG000684 | Rev A, 10x genomics”, followed by the “Visium HD Spatial Gene Expression Reagent Kits User Guide, CG000685 | Rev B, 10x genomics”. For scRNAseq, the libraries were sequenced with a HiSeqX Illumina sequencer at a depth of 50 000 reads per cell (Macrogen), wile for Visium HD the libraries were then sequenced with a NextSeq 2000 P2 Illumina sequencer at a depth of 400 million reads (Biomics Plateform, Pasteur Institute).

### ScRNA-seq and Visium HD analysis

For scRNA-seq, raw sequencing data were demultiplexed using bcl2fastq conversion software (Illumina), processed using the Cell Ranger pipeline (version 7.1.0, 10x genomics) and analyzed with Partek Flow analysis software (Partek, St. Louis, Missouri). The following criteria for the selection of healthy cells and exclusion of duplicates were applied: 500-6000 genes, 1250-25000 counts, 0-8% mitochondrial reads and 0-10% ribosomal reads.

For Visium HD, raw sequencing data were demultiplexed using bcl2fastq conversion software (Illumina), processed using the Space Ranger pipeline (version 3.1.2, 10x genomics) and analyzed with Partek Flow analysis software (Partek, St. Louis, Missouri). A binning of 8 µm² (instead of 2 and 16) was considered, to reach a single-cell resolution and a sufficient number of reads per bin to perform differential gene expression analysis. The bins not overlapping the tissue sections were filtered out by increasing the quality threshold for minimum number of genes expressed per bin, varying between samples.

For both datasets, gene expression was normalized in counts per million (by dividing by the total number of mapped reads per sample and multiplying by 1 × 10^6^). A count of 1 was then added before log_2_ transformation of the data. Genes not expressed in 99.9% of cells were filtered out. Cells were classified into cell- types with automated graph-based clustering. If necessary, the bins/cells were reassigned manually based the combined expression patterns of canonical markers. The classification was validated by comparing the top 100 most differentially expressed genes between these cell types with published scRNA-seq datasets from the mouse cochlea. For Visium HD data, the classification was additionally supported by comparing the assigned cell-types to the cell-types observed onto the histological tissue sections at the corresponding coordinates.

### Statistical and data analysis

Principal component analysis focusing on the most variable features was performed and the top 15 principal components selected before data visualization with a uniform manifold approximation and projection (UMAP) using Cosine distance metric. Automatic graph-based clustering was performed using Louvain algorithm and Euclidean distance metric The most differentially expressed genes were identified by Student’s t-tests on the classified cell types, comparing each group, one-by-one, with all the other groups together. The upregulated genes were identified as markers, and were sorted on the basis of their p-values. Differential gene expression analysis was performed using the Hurdle model, where p-values were corrected for multiple comparisons with FDR-step-up. For heat map/hierarchical clustering, the average linkage method was used to determine cluster distance metrics and the Euclidean method was used to determine point distance metrics. For bubble plot analysis, the mean gene expression level and the percent of cells with a gene expression value above 0 were considered.

---

**Fig. S1.**
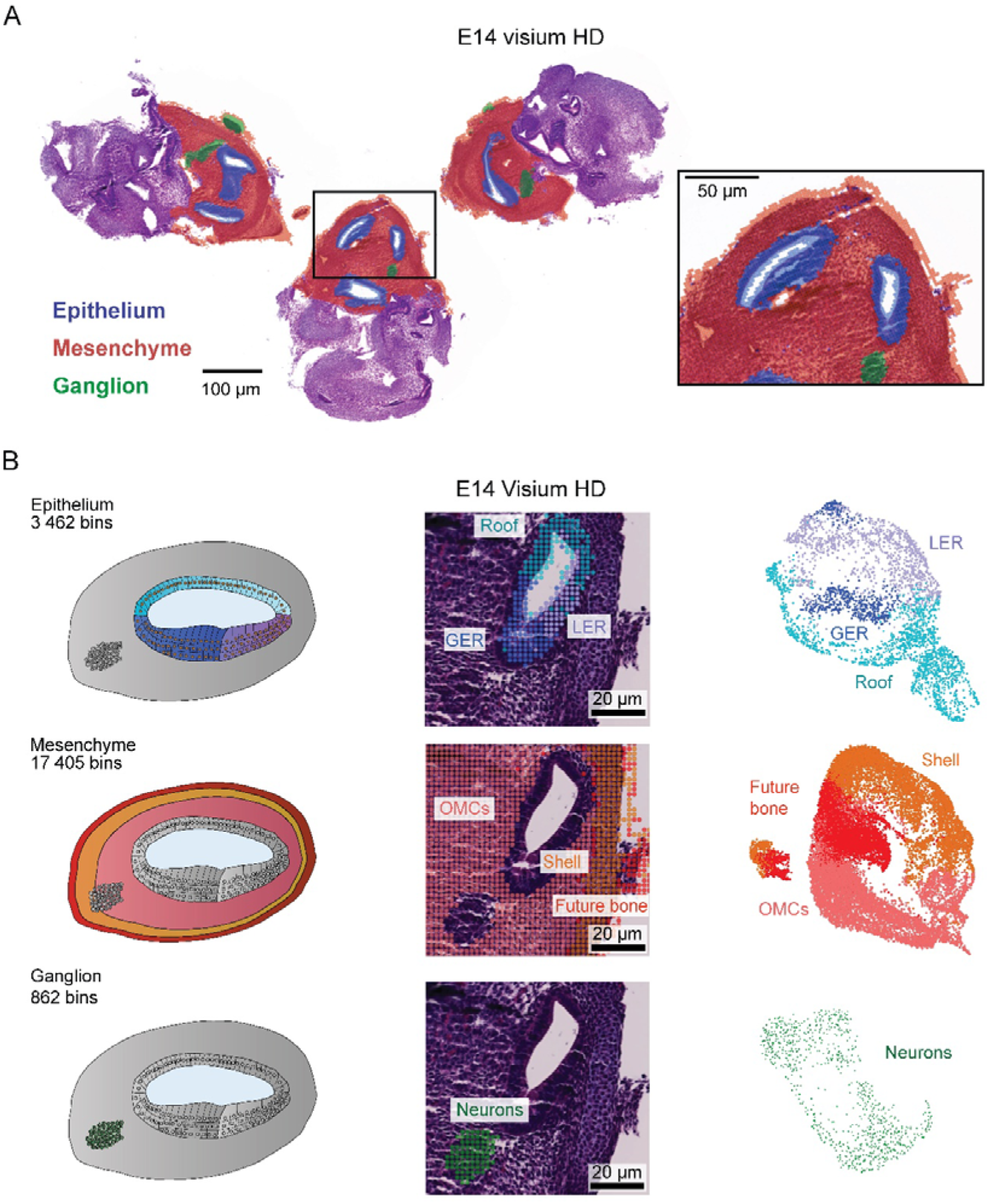
Cell-type assignment in Visium HD datasets on E14. (A) Histological FFPE cross sections of the three E14 analyzed cochleae stained with H&E and overlayed with the filtered bins color coded according to their main cochlear ensemble. (B) Left: Cross-section schematic of a cochlear turn where the cell-types (their markers indicated in parenthesis) from either the epithelium (Roof (Oc90), GER (Tecta, Tectb, Clic6), LER (Fst), mesenchyme (OMCs (Pou3f4, Tbx18), Shell (Acan), Future bone) or ganglion (neurons (Pvalb, Scng, Celf3)) are color coded. Middle: Histological cross sections from FFPE samples stained with H&E, overlayed with the filtered bins following the same color code. Right: UMAP Plots of the total number of bins analyzed in each cochlear ensemble following the same color code.

**Fig. S2.**
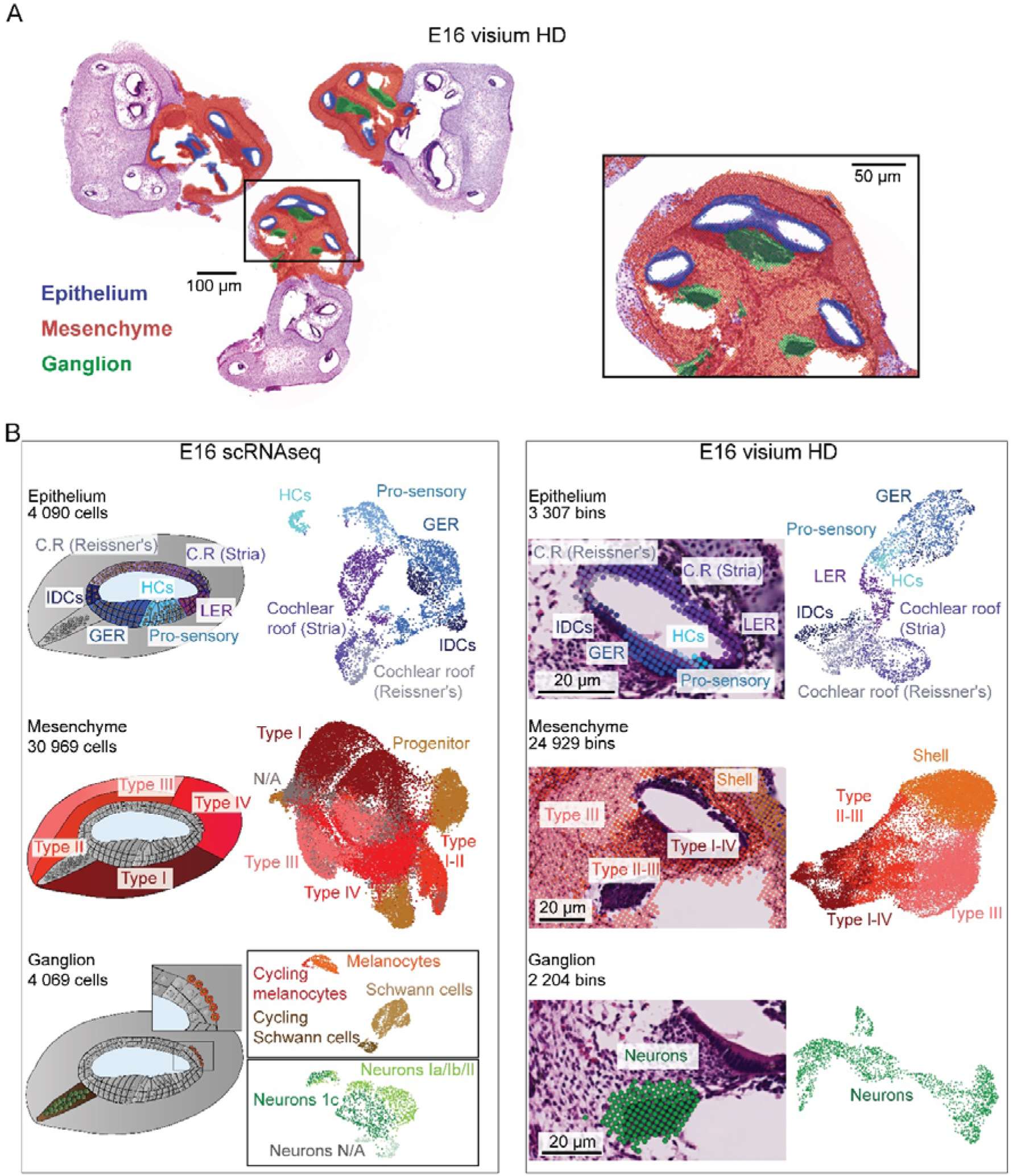
Assignment of cell-types in Visium HD datasets on E16. (A) Histological FFPE cross sections of the three E16 analyzed cochleae stained with H&E and overlayed with the filtered bins color-coded according to their main cochlear ensemble (B) Left: scRNA-seq dataset with diagrams of a cochlear turn cross-section with the corresponding UMAP plots of the total number of cells analyzed in each cochlear ensemble where the cell-types, with their markers indicated in parenthesis, from either the epithelium (Roof Stria (Oc90, Sparcl1), Roof Reissner (Oc90, Otoa), interdental cells (IDCs) (Otoa), GER (Calb1, Tecta, Clic5), HCs (Atoh1, Pou4f3), pro-sensory (Tectb, Sox2, Ntf3), LER (Bmp4, Fst)), mesenchyme (progenitors (Pou5f1, Nanog, Mki67), type I-II (Aldh1a2, Kcnk2), type-III (Penk, Col1a1, Ibsp), type I-IV (Coch, Car3, Otor)) or ganglion (Schwann cells (Mpz, Prx), Melanocytes (Dct, Mlana, Tyr), neurons (Nefm, Nefl, Scrt2), Ia/Ib/II (Prph, Islr2) and Ic (Pou4f1, Lypd1)) are color-coded. Right: Visium HD dataset with histological cross sections from FFPE samples stained with H&E, overlayed with the filtered bins, alongside with the UMAP plots of the total number of bins analyzed in each cochlear ensemble following the same color code as for scRNA-seq.

**Fig. S3.**
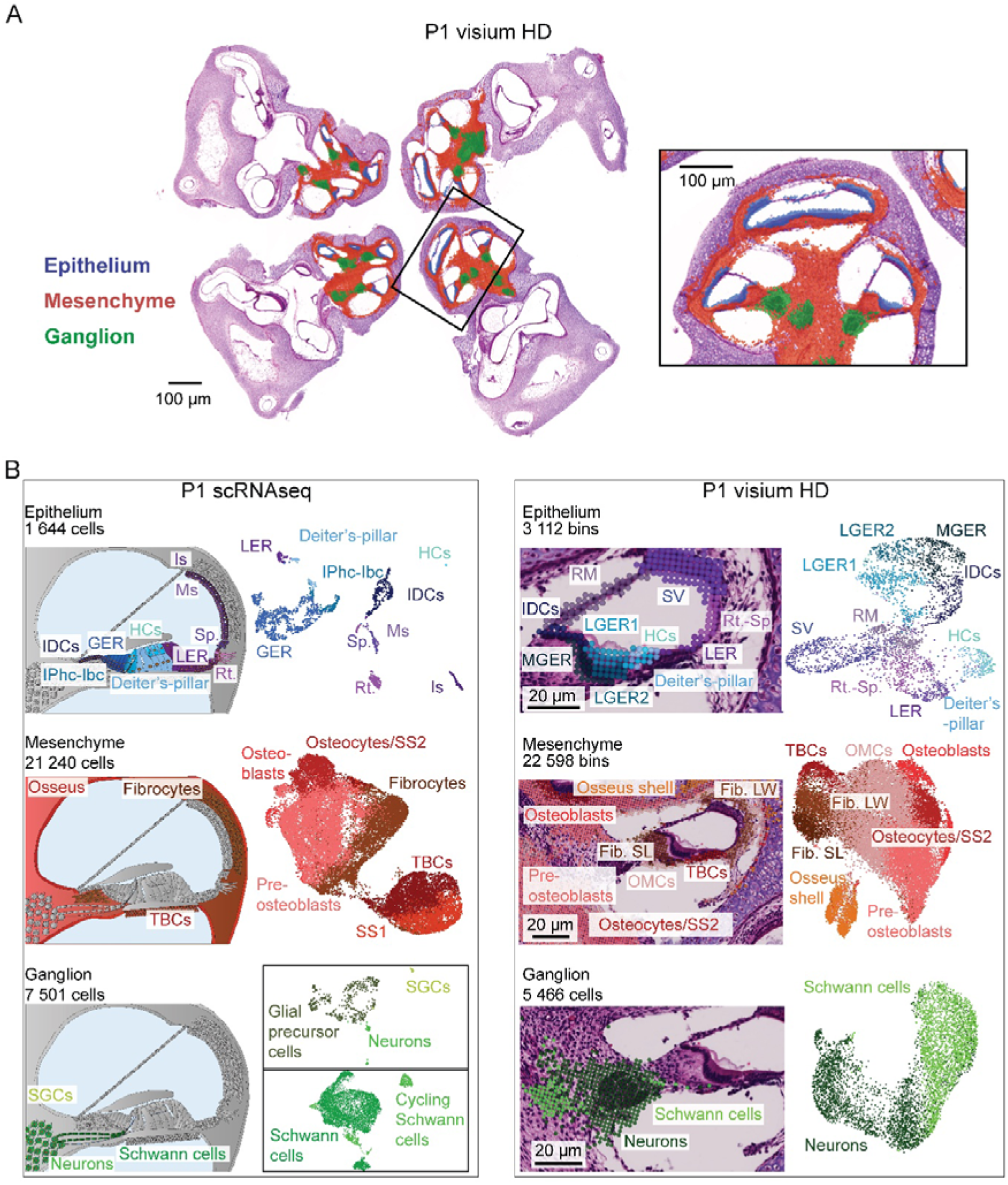
Cell-type assignment in the P1 Visium HD dataset. (A) Histological FFPE cross sections of the four P1 analyzed cochleae stained with H&E, overlayed with the filtered bins color-coded according to their main cochlear ensemble. (B) Left: diagrams illustrating the different cell types, with their markers indicated in parenthesis, of the cochlea with the corresponding UMAP plots for each cochlear ensemble where are color-coded the cell-types from either the epithelium (interdental cells (IDCs) (Otoa), GER (LGER1/2 & MGER cells) (Calb1, Tecta, Crabp5), inner phalangeal-border cells (Ano1, Cpa6, Slitrk6), HCs (Atoh1, Pou4f3), Deiter’s and pillar cells (Cep41, Prox1, Fgfr3), LER (Rubie, Emx2), Reissner membrane cells (RM) (Slc26a7, Vmo1, Tmem72 and Lrp2), marginal stria cells (Ms) (Kcne1, Kcnq1, Bsnd), intermediate stria cells (Is) (Dct, Tyr, Mlana), root cells (Rt.) (Slc26a4, Epyc), spindle cells (Sp.) (Slc26a4, Anxa1), mesenchyme (pre-osteoblasts (Dlx5, Runx2, Gdf10), osteoblasts (Bglap, Ifitm5, Ibsp), osteocytes (Hpca, Gcg), TBCs (Notum, Emilin2), fibrocytes (Car3, Otos, Otor) or ganglion (Schwann cells (Mpz, Prx), Satellite glial cells (Mog, Mag, Ermn), glial precursor cells (Lhfpl3, Olig1, Olig2), neurons (Tubb3, Snap25, Neurod1)). Right: Visium HD dataset with histological cross sections from FFPE samples stained with H&E, overlayed with the filtered bins, alongside with the UMAP plots of the total number of bins analyzed in each cochlear ensemble following the same color code as for scRNA-seq.

**Fig. S4.**
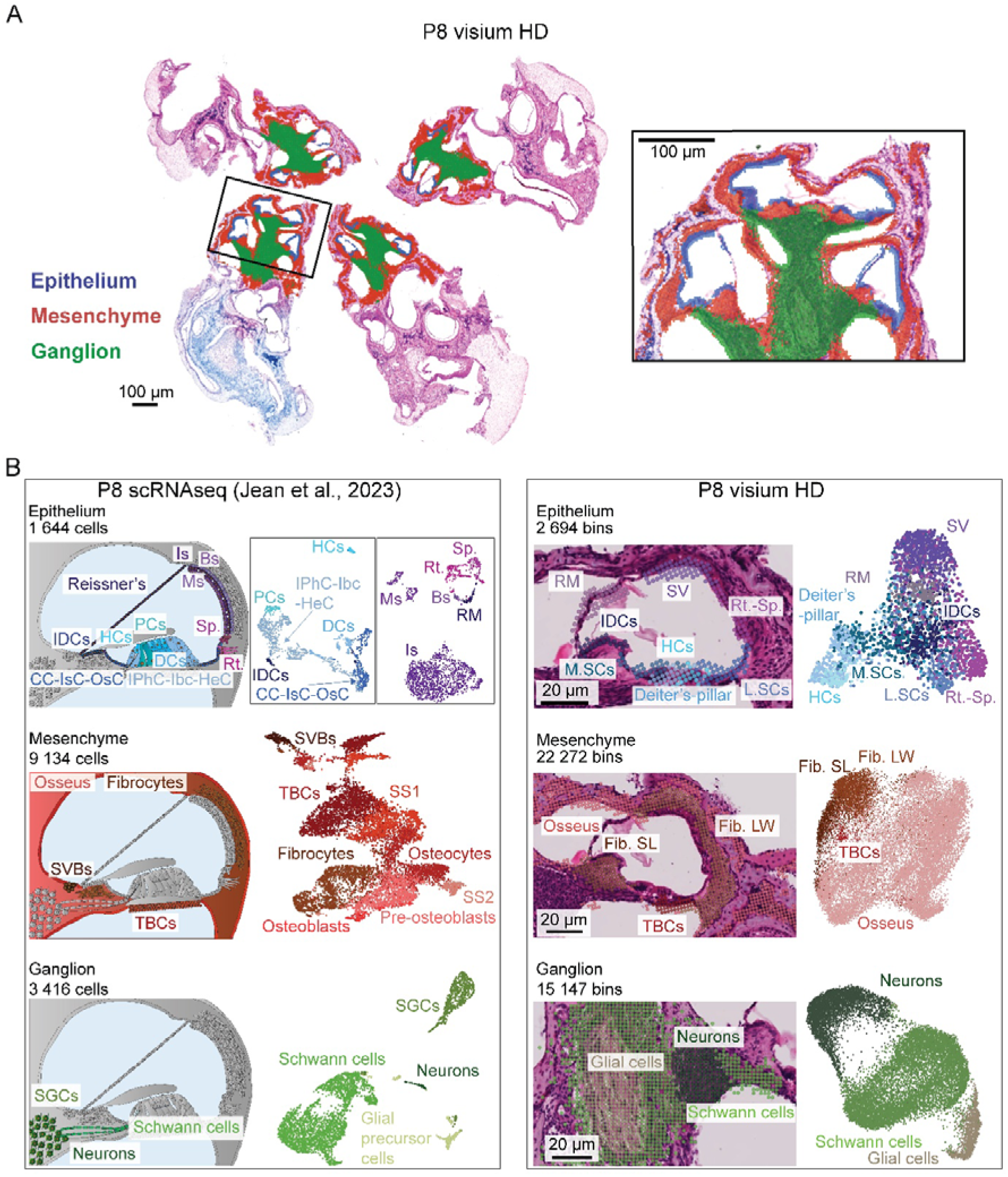
Cell-type assignment in Visium HD datasets on P8. (A) Histological FFPE cross sections of the four P8 analyzed cochleae stained with H&E, overlayed with the filtered bins color coded according to their main cochlear ensemble. (B) Left: diagrams illustrating the different cell types of the cochlea with the corresponding UMAP plots from the published P8-P12-P20 scRNA-seq dataset (Jean et al., 2023) for each cochlear ensemble where are color-coded the cell-types, with their markers indicated in parenthesis, from either the epithelium (interdental cells (IDCs) (Otoa), medial supporting cells (M.SCs) (Epyc, Tectb), HCs (Otof, Slc26a5), Deiter’s and pillar cells (Cep41, Ush1c, Tectb), lateral supporting cells (L.SCs) (Epyc, Bmp4), Reissner membrane cells (RM) (Slc26a7, Vmo1, Tmem72 and Lrp2), marginal stria cells (Ms) (Kcne1, Kcnq1, Bsnd), intermediate stria cells (Is) (Dct, Tyr, Mlana), fibrocytes (Car3, Otos, Otor)) or ganglion (Schwann cells (Mpz, Prx), Satellite glial cells (Mog, Mag, Ermn), neurons (Tubb3, Snap25, Neurod1)). Right: P8 Visium HD dataset with histological cross sections from FFPE samples stained with H&E, overlayed with the filtered bins, alongside with the UMAP plots of the total number of bins analyzed in each cochlear ensemble following the same color code as for scRNA-seq.

**Fig. S5.**
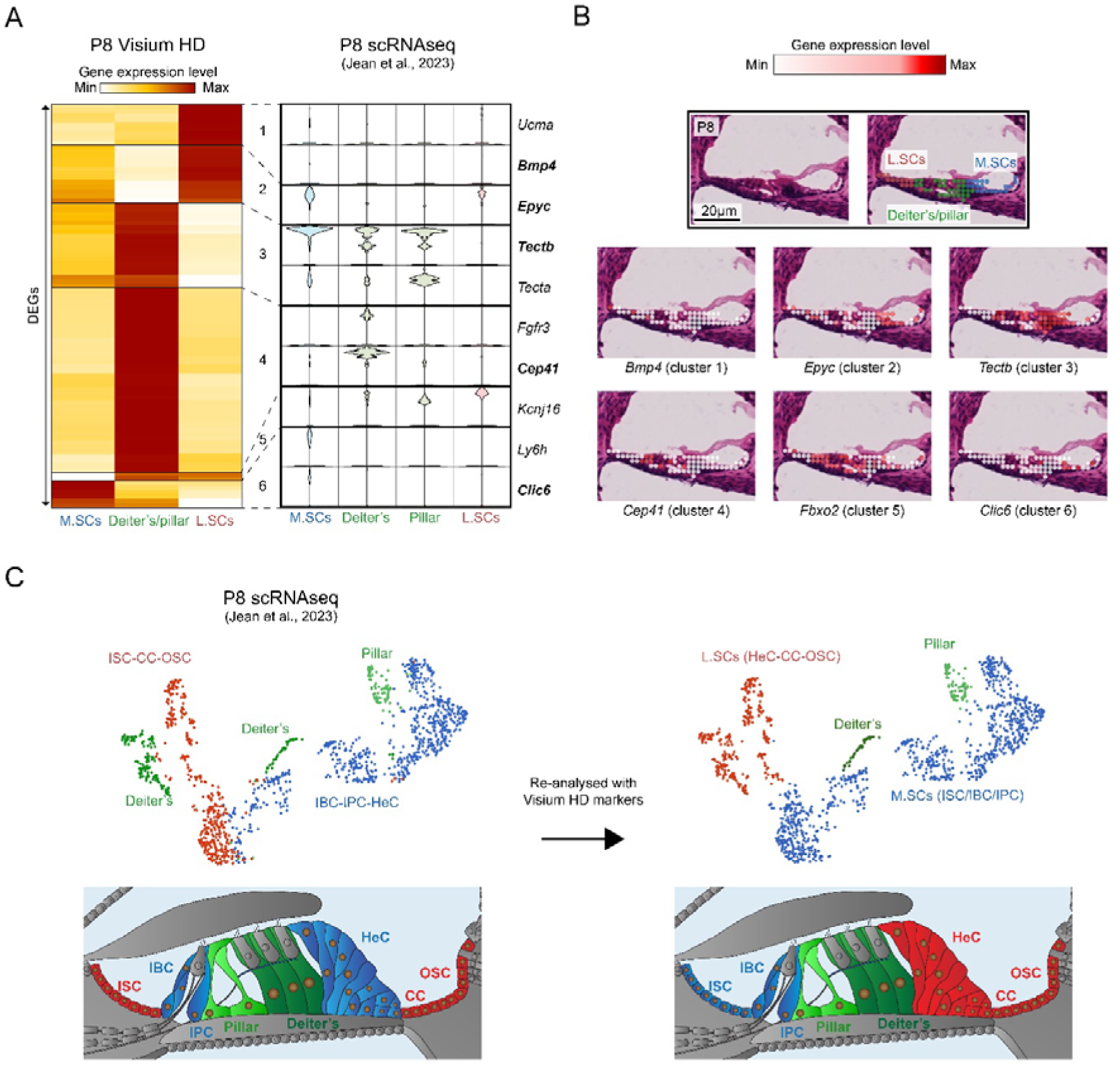
Radial differential gene expression analysis between supporting cell subtypes on P8. (A) Left: Heat map/hierarchical clustering of the DEGs distributed between the GER, Deiter’s-pillar cells and LER on P8. Right: Violin plots depicting the expression of the DEGs found in the Visium HD dataset, in the scRNAseq dataset at the same age (Jean et al., 2023). (B) Histological Visium HD cross sections from P8 FFPE samples stained with H&E, and overlayed with the filtered bins color-coded according to the gene expression level. (C) Left: UMAP plot of the SCs in our published P8 scRNA-seq (Jean et al., 2023), with the corresponding diagram illustrating a cross section of the cochlea where the SCs are color-coded accordingly. Right: The data has been reanalyzed using the markers obtained from the Visium HD dataset.

**Fig. S6.**
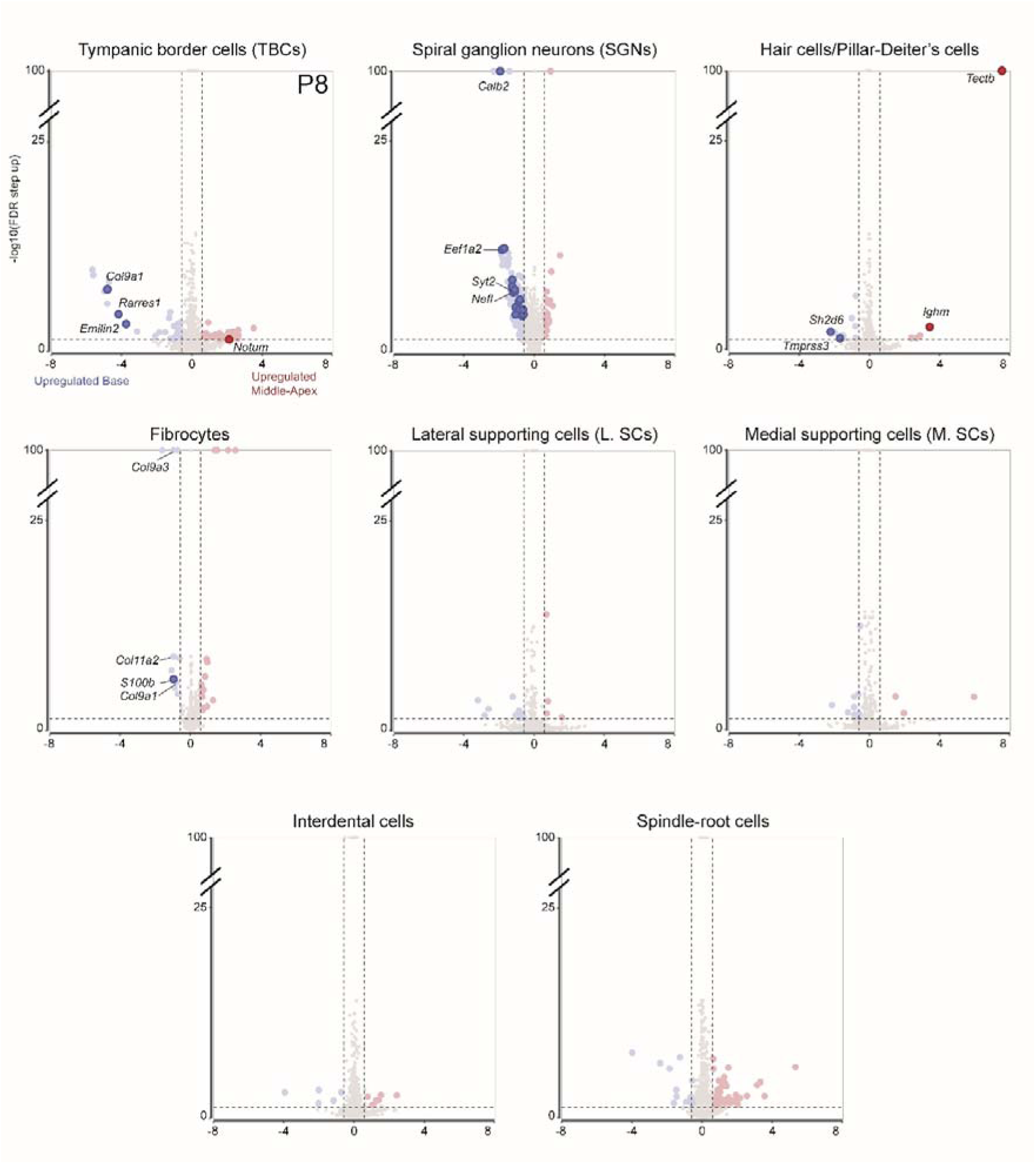
Tonotopic differential gene expression analysis on P8. Volcano plots displaying the differential pattern of gene expression between apical (in red) and basal (in blue) cells for the different cochlear cell types on P8. Fold-changes in expression are indicated on the x axis, with positive values indicating an upregulation at the middle-apex and negative values indicating an upregulation at the base. The false-discovery rate-corrected p-value is indicated on the y axis. Only genes with a fold-change in expression of at least 1.5 in either direction (-1.5/+1.5) and an FDR-corrected p-value less than 0.05 are colored. The genes following a strict tonotopic gradient (A>MA>MB>B or B>MB>MA>A) are highlighted. The genes with their names written are the ones illustrated in the main figure.

**Fig. S7.**
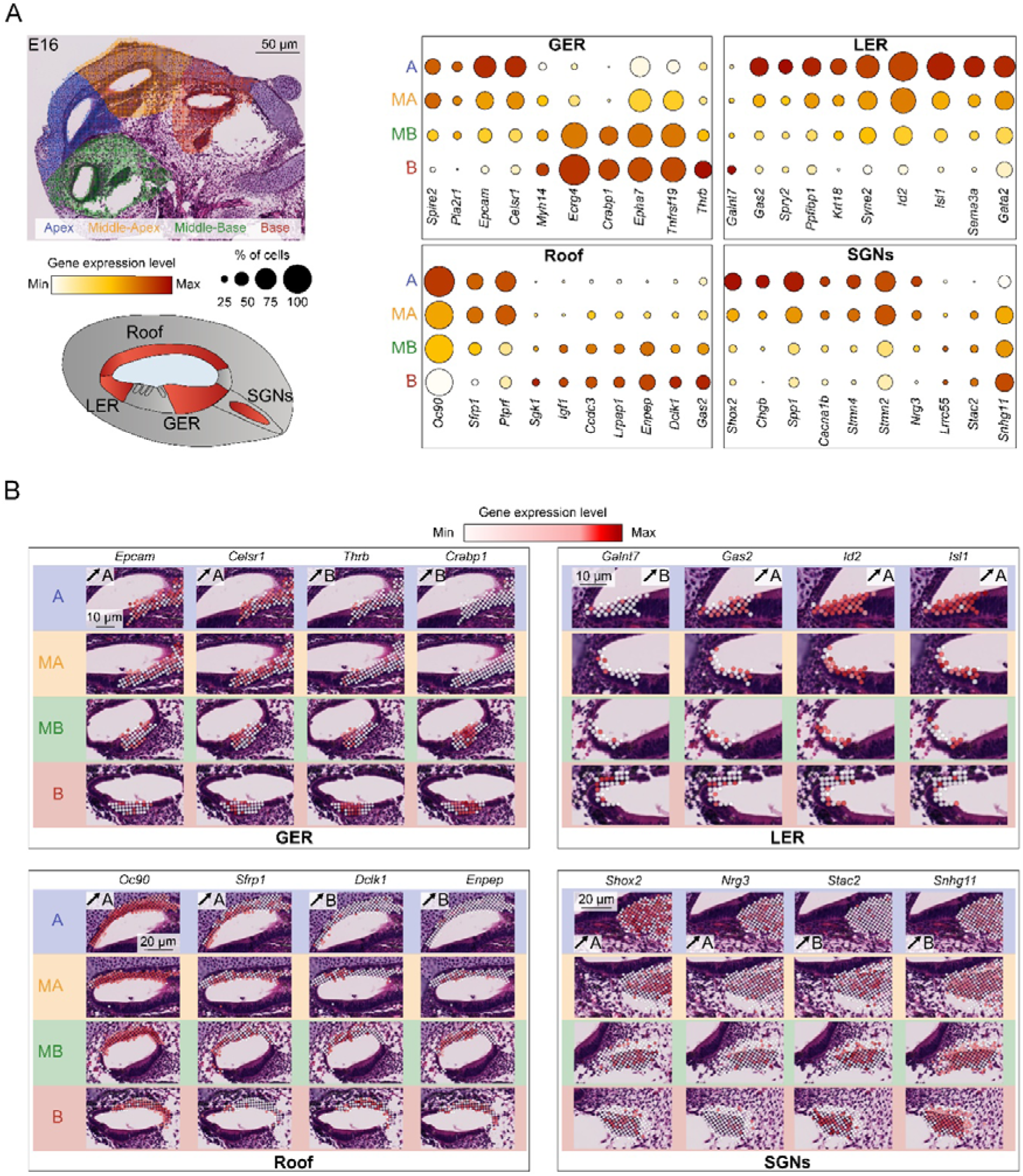
Tonotopic gradient of gene expression in the GER, LER, and Roof cells as well as SGNs on E16. (A) Top left: Histological Visium HD cross section from FFPE samples stained with H&E, showing the cochlear tissue analyzed, overlayed with the filtered bins color-coded according to their tonotopic location at the apex (in blue), middle-apex (in yellow), middle-base (in green) and the base (in red). Bottom left: Diagram illustrating the four cell types of the cochlea analyzed. Right: Bubble plot analysis comparing the expression level of a subset of genes having a tonotopic gradient of expression either from base to apex, or from apex to base. The gene expression level is color-coded while the size of the spots indicates the proportion of cells expressing the genes. (B) Histological Visium HD cross sections from FFPE samples stained with H&E, overlayed with the filtered bins color-coded according to the expression of the gene of interest, at the different tonotopic localizations for each of the analyzed cell-types.

**Fig. S8.**
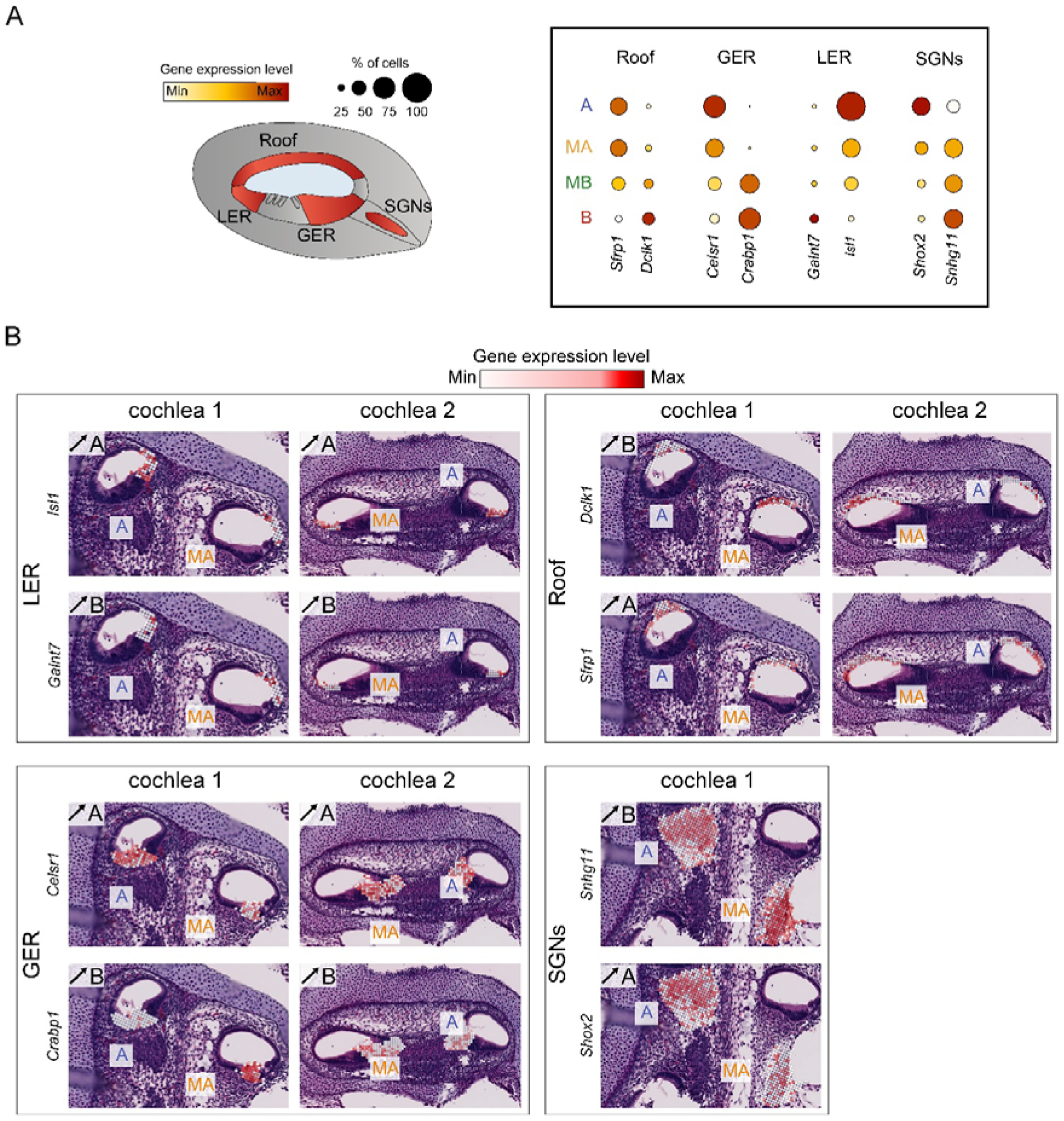
Tonotopic gradient of gene expression in the GER, LER and Roof cells as well as SGNs on E16. (A) Left: Diagram illustrating the four cochlear cell-types analyzed. Right: Bubble plot analysis comparing the expression level of a subset of genes having a tonotopic gradient of expression either from base to apex, or from apex to base. The gene expression level is color-coded while the size of the spots indicates the proportion of cells expressing the genes. (B) Histological Visium HD cross sections from FFPE samples stained with H&E, overlayed with the filtered bins color-coded according to the expression of the gene of interest, at the different tonotopic localization for each of the analyzed cell-types.

**Fig. S9.**
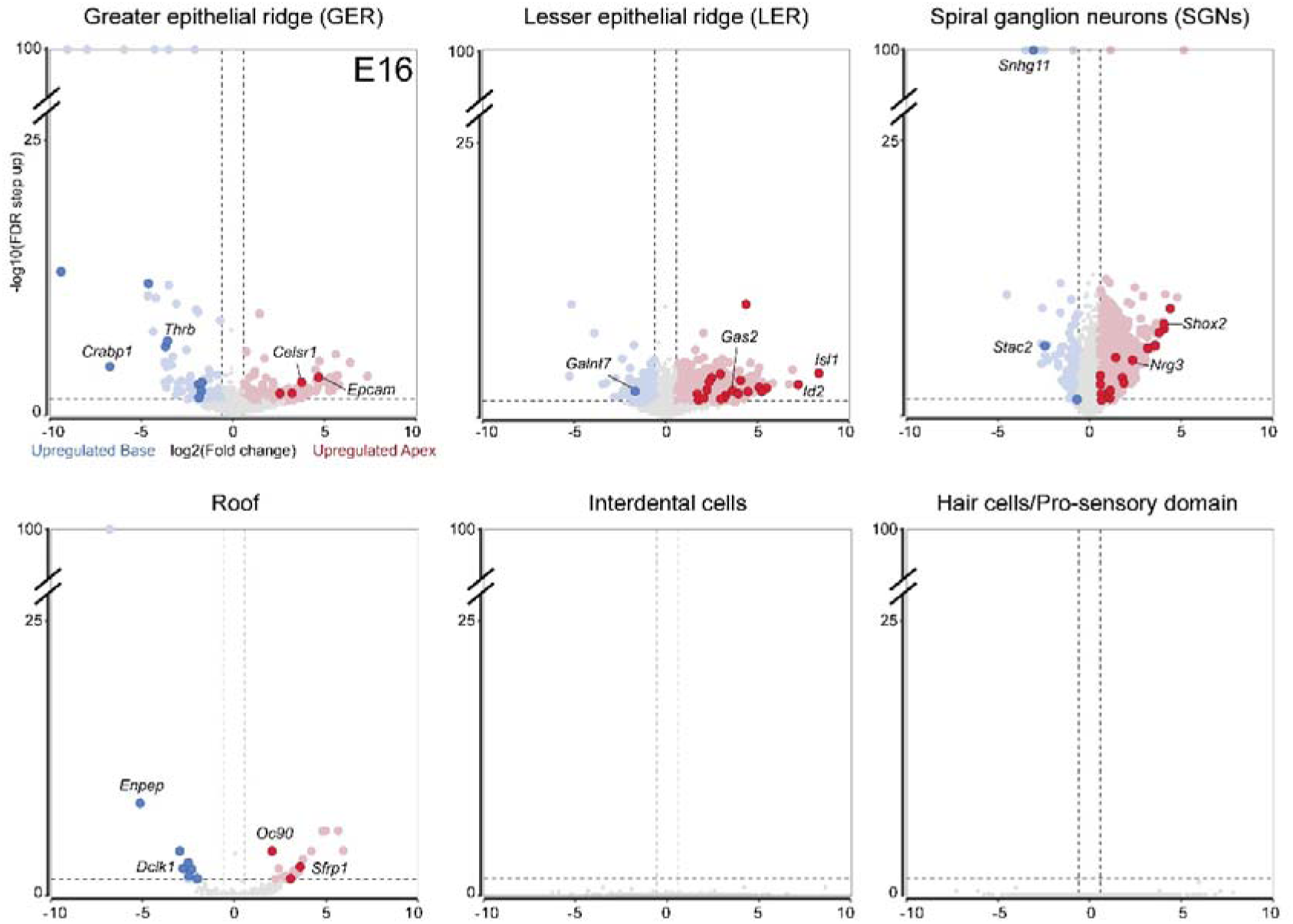
Differential gene expression between apical and basal cochlear cells on E16. Volcano plots displaying DEGs between the apex (in red) and base (in blue) of the cochlea for the different cochlear cell types on E16. Fold-changes in expression level are indicated on the x axis, with positive values indicating an upregulation at the apex and negative values indicating an upregulation at the base. The false-discovery rate-corrected p-value is indicated on the y axis. Only genes with a fold-change in expression of at least 1.5 in either direction (-1.5/+1.5) and an FDR-corrected p-value less than 0.05 are colored. The genes following a strict tonotopic gradient (A>MA>MB>B or A<MA<MB<B) are highlighted. The genes with their names written are the ones illustrated in the main figure.

**Fig. S10.**
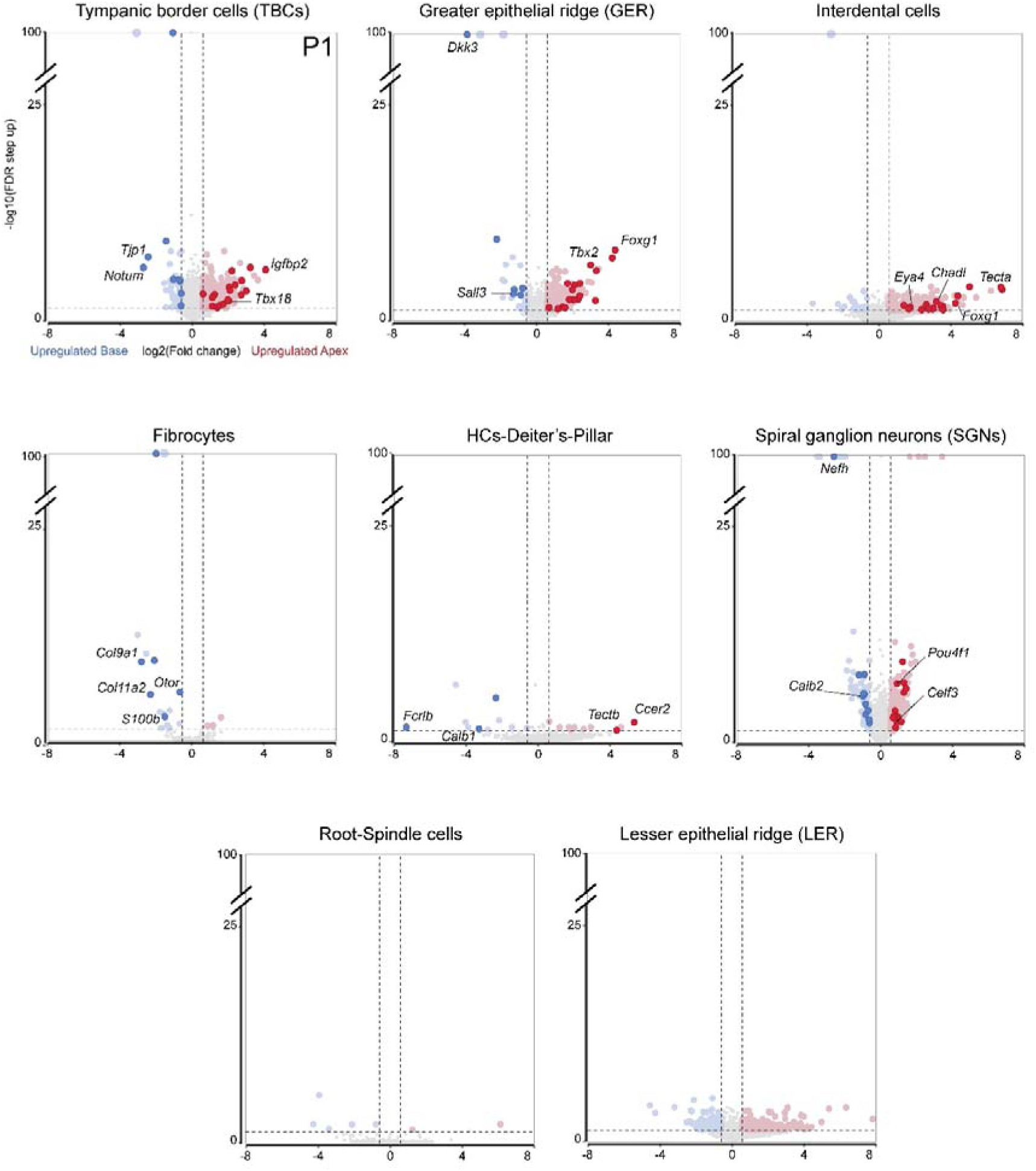
Differential gene expression apical and basal cells on P1. Volcano plots displaying DEGs between the apex (in red) and base (in blue) of the cochlea for the different cochlear cell types on P1, and between middle-apex and base for the spiral ganglion neurons, absent at the apex, Fold-changes in expression are indicated on the x axis, with positive values indicating an upregulation at the apex and negative values indicating an upregulation at the base. The false-discovery rate-corrected p-value is indicated on the y axis. Only genes with a fold-change in expression of at least 1.5 in either direction (-1.5/+1.5) and an FDR-corrected p-value less than 0.05 are colored. The genes following a strict tonotopic gradient (A>MA>MB>B or B>MB>MA>A) are highlighted. The genes with their names written are the ones illustrated in the main figure.

**Fig. S11.**
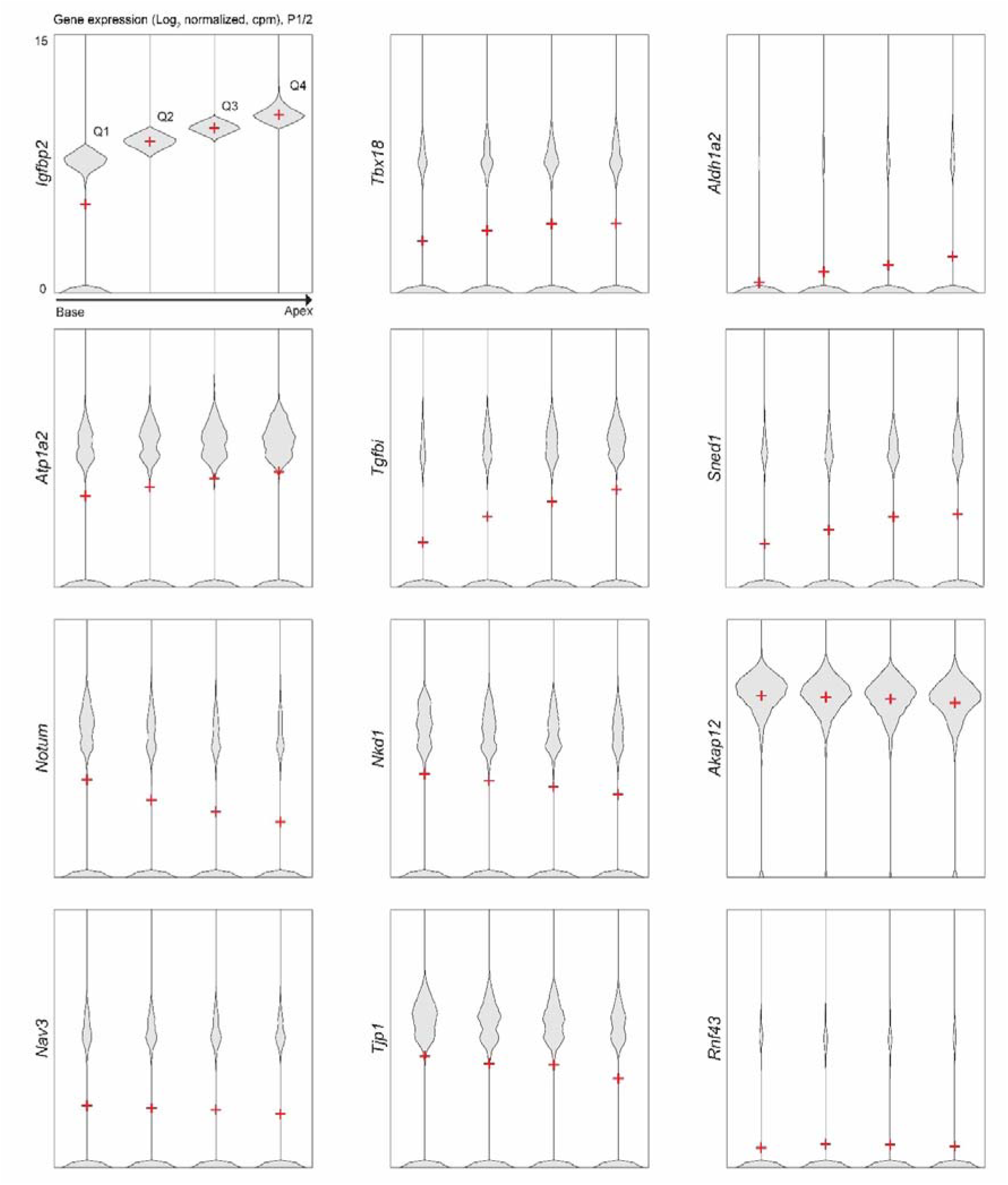
DEGs along the tonotopic axis in TBCs of the P1 scRNA-seq dataset. Violin plots showing the levels of expression on P1 (counts per million, log_2_-normalized) of genes differentially expressed in tympanic border cells and classified into four equal quartiles (Q1 to Q4) assumed to represent four tonotopic subregions of the cochlea from the base to the apex according to Igfbp2 expression. Means are represented by a red cross.

**Fig. S12.**
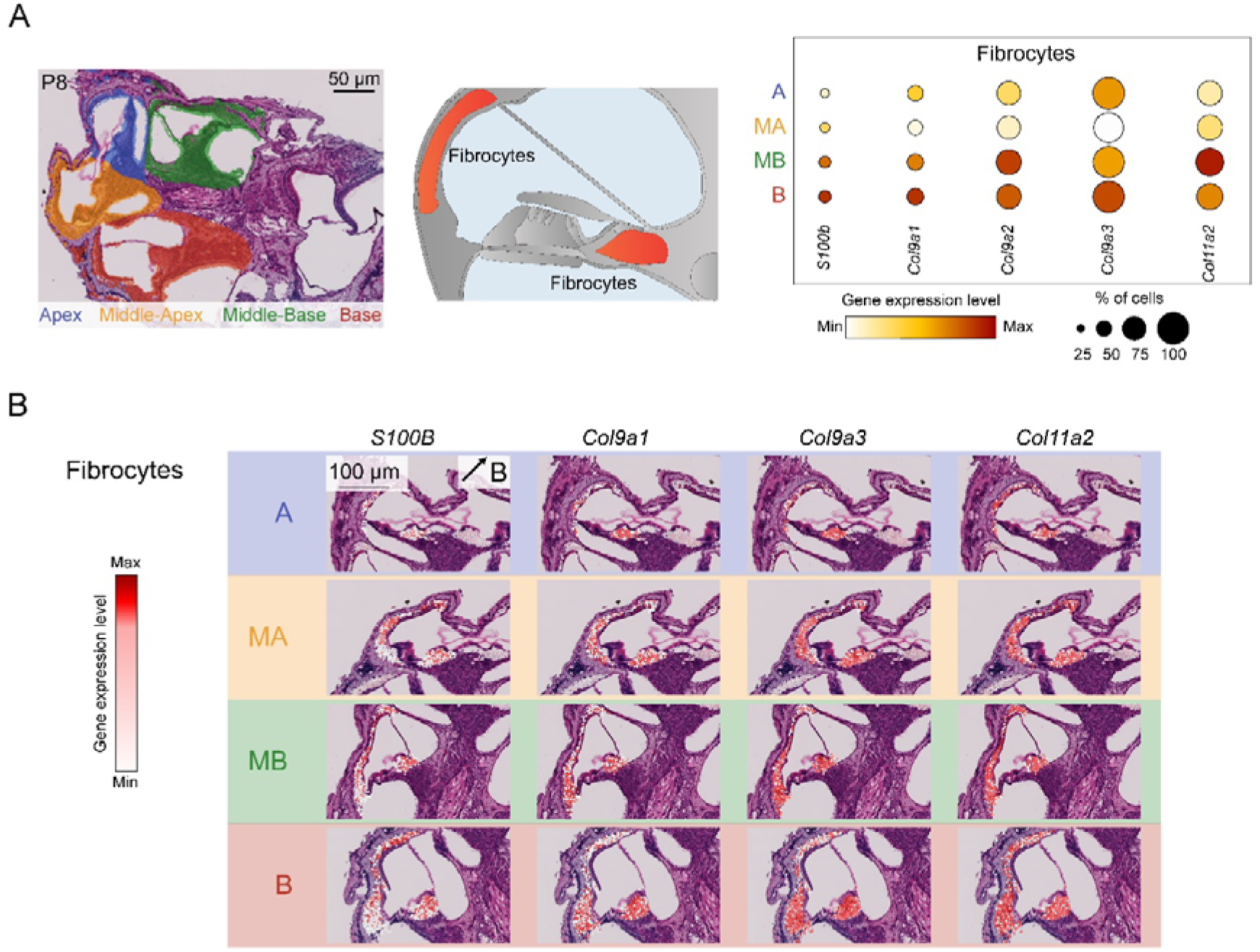
DEGs along the tonotopic axis in fibrocytes on P8. (A) Left: Histological Visium HD cross sections from FFPE samples stained with H&E, showing a cochlea analyzed, overlayed with the filtered bins color-coded according to their tonotopic location being the apex (in blue), middle-apex (in yellow), middle-base (in green) and the base (in red) of the cochlea. Middle: Diagram illustrating the cochlear cell-types analyzed. Right: Bubble plot analysis comparing the expression level of a subset of genes presenting a tonotopic gradient of expression either from base to apex, or from apex to base. The gene expression level is color-coded from white (minimum) to red (maximum), while the size of the spots indicates the proportion of cells expressing the genes. (B) Histological cross sections from FFPE samples stained with H&E, overlayed with the filtered bins color-coded according to the gene expression, at the different tonotopic localizations.

**Fig. S13.**
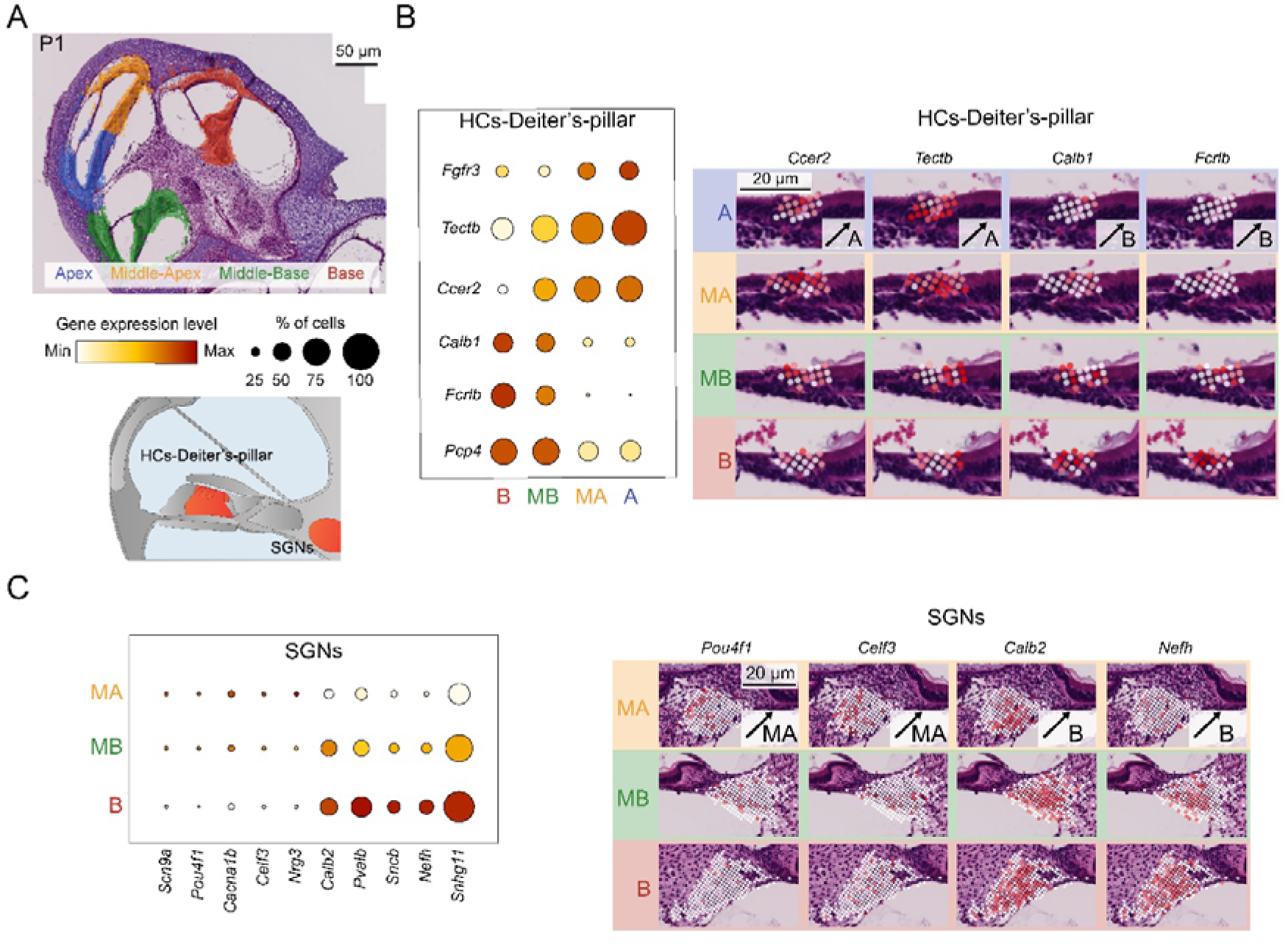
DEGs along the tonotopic axis in HCs-Deiter’s-pillar cells and SGNs on P1. (A) Top left: Histological Visium HD cross sections from FFPE samples stained with H&E, showing a cochlea analyzed, overlayed with the filtered bins color-coded according to their tonotopic location being the apex in blue, middle-apex in yellow, middle-base in green and the base in red. Bottom left: diagram illustrating the analyzed cell-types. (B) Left: Bubble plot analysis comparing the expression level of a subset of genes presenting a tonotopic gradient of expression either from base to apex, or from apex to base. The gene expression level is color-coded, while the size of the spots indicates the proportion of cells expressing the genes. Right: histological Visium HD cross sections from FFPE samples stained with H&E, overlayed with the filtered bins color-coded according to the gene expression, at the different tonotopic localizations.

